# Identification of the protein precursor for thyroid hormone synthesis in the basal chordate ascidian *Styela clava*

**DOI:** 10.64898/2025.12.08.693098

**Authors:** Jin Zhang, Likun Yang, Brice Beinsteiner, Yun Ma, Jiankai Wei, Haiyan Yu, Vincent Laudet, Bo Dong

**Author notes:** These authors contribute equally to this work (Jin Zhang, Likun Yang). Correspondence (B.D.).

## Abstract

Thyroid hormones (THs) play crucial roles in development, growth, and metabolism across the animal kingdom^1,2^. Although the mechanism of TH synthesis in vertebrates has been well described, it remains elusive in invertebrates due to the lack of identified orthologs of the thyroglobulin (TG), the protein precursor for TH synthesis^3,4^. Here, we identified a functional TG protein encompassing multiple TG type 1 (Tg1)-domains, encoded by the *ScTg-like* gene in proto-vertebrate ascidian *Styela clava* based on the immunoprecipitation-mass spectrometry combined with phylogenetic and expression pattern analyses. *In vitro* iodination experiments demonstrated that ScTG-like has the capacity to provide hormonogenic sites for TH synthesis. Furthermore, *in vivo* gene knockdown of *ScTg-like* revealed that the larval metamorphosis was significantly inhibited as a consequence of diminished TH levels, indicating that *ScTg-like* is responsible for the production of endogenous TH. Additionally, an invaginated follicle-like structure was first discovered at the central anterior of the larval trunk where both ScTG-like protein and THs were deposited, suggesting that this structure serves as the location for THs synthesis and storage, functionally similar to vertebrate thyroid follicles. Moreover, we analyzed the structural features of ScTG-like protein and detected the similarly organized Tg1-containing proteins in multiple bilaterian phyla, suggesting that endogenous TH synthesis may be an ancestral, possibly synapomorphic, trait of bilaterians. Our study is the first to identify the TH protein precursor outside vertebrates by providing the direct evidence of endogenous *de novo* TH synthesis, which offers insights into the evolution of TG protein and the emergence of a major vertebrate endocrine system.

## Main

In vertebrates, thyroid hormone (TH) biosynthesis involves many essential proteins, the functions of which have been well described^5,6^. Among them, thyroglobulin (TG) is the protein precursor of TH, serving as the highly specialized matrix for iodination of specific tyrosine residues to form monoiodotyrosine (MIT) and diiodotyrosine (DIT), and for their coupling to generate triiodothyronine (T3) or thyroxine (T4)^5,6^. In all vertebrates, TG is multi-domain protein with a high-molecular-weight (approximately 330 kDa)^5^. It consists of a signal peptide, three types of TG type domains (Tg1-3) repeated several times and separated by different linkers and one cholinesterase-like (ChEL) domain in C-terminus^7^. The Tg domains are essential for TG functions including proper folding, iodination, iodine storage and release^8,9^. The ChEL domain is implicated in TG maturation and trafficking, as well as the *de novo* synthesis of T3^10–12^. TG proteins, in terms of their domain distribution, structure, and key hormonogenic sites, are highly conserved in vertebrates^4^. The current view of TG evolution holds that lampreys may be the most ancient vertebrate species with a TG protein^13,14^, since the orthologs of TG have not been identified outside of vertebrates.

Outside vertebrates, THs are detectable and play crucial functions in development, especially metamorphosis^15–17^. Yet, how invertebrates synthesize THs remains unclear, as they lack vertebrate-like TG proteins and no functional precursor has been convincingly identified. Previous studies have reported that several marine organisms harbor the ability to synthesize TH *in vivo*. The sea urchin and sea hare have ability to incorporate iodine in T4 formation^18^. In amphioxus, the iodoprotein(s) have been described in endostyle, which exhibit the similar sedimentation coefficient as vertebrate TG and are capable of producing THs^19^. In ascidian, the iodoprotein(s), with similar sedimentation coefficient to vertebrate TG, are mainly distributed in endostyle, pharynx, and tunic^20–23^. Endostyle is a specialized pharyngeal organ, with zone 7-9 representing the thyroid-equivalent region in ascidian adult^22,24^. Antibody against mammalian TG could specifically label the thyroid-equivalent region in ascidian *Styela clava* endostyle^25^, implying that the immunoreactive protein(s) may be ascidian TG-like protein(s) potentially involved in TH synthesis. However, the exact sequences of these potential iodoproteins or TG-like protein(s) and their functions in the developmental physiology have remained unexplored.

### Identification of *S. clava* Tg1 domain-containing proteins

To determine whether ascidian process vertebrate-like TG protein, we prepared an antibody against mammalian (*Bos taurus*) TG protein (anti-*Bt*TG) and performed immunofluorescence staining on *S. clava* endostyle. Consistent with the previous results^25^, the antibody specifically labeled the thyroid-equivalent region of endostyle (Fig. 1a-b), suggesting that the TG-like protein(s) may be present in *S. clava*. To further identify the immunolabeled protein(s), immunoprecipitation coupled with mass spectrometry (IP-MS) was conducted to precipitate the immunolabeled protein(s) within the endostyle tissue lysates (Supplementary Fig. 1a). A total of ten Tg1 domain-containing proteins were identified from IP-MS data, including Sc.CG.S33.65, Sc.CG.S284.31, Sc.CG.S566.2, Sc.CG.S166.41, Sc.CG.S140.18, Sc.CG.S6.11, Sc.CG.S3.11, Sc.CG.S79.69, Sc.CG.S131.45, and Sc.CG.S55.56, which represent five-sixths of all predicted Tg1 domain-containing proteins in *S. clava* genome (Fig. 1c, Supplementary Fig. 1b-e). Furthermore, we performed a phylogenetic analysis comparing these Tg1 domain-containing proteins with vertebrate TG proteins, Tg1 domain-containing proteins in human genome^26^ and predicted TG-like protein in amphioxus and sea urchin^9^. The result revealed that Sc.CG.S284.31 and Sc.CG.S55.56 proteins clustered within the clade of vertebrate TG proteins, suggesting that they were the closest relatives to vertebrate TG proteins among all the identified Tg1 domain-containing proteins (Fig. 1d-e).

**Figure 1.**
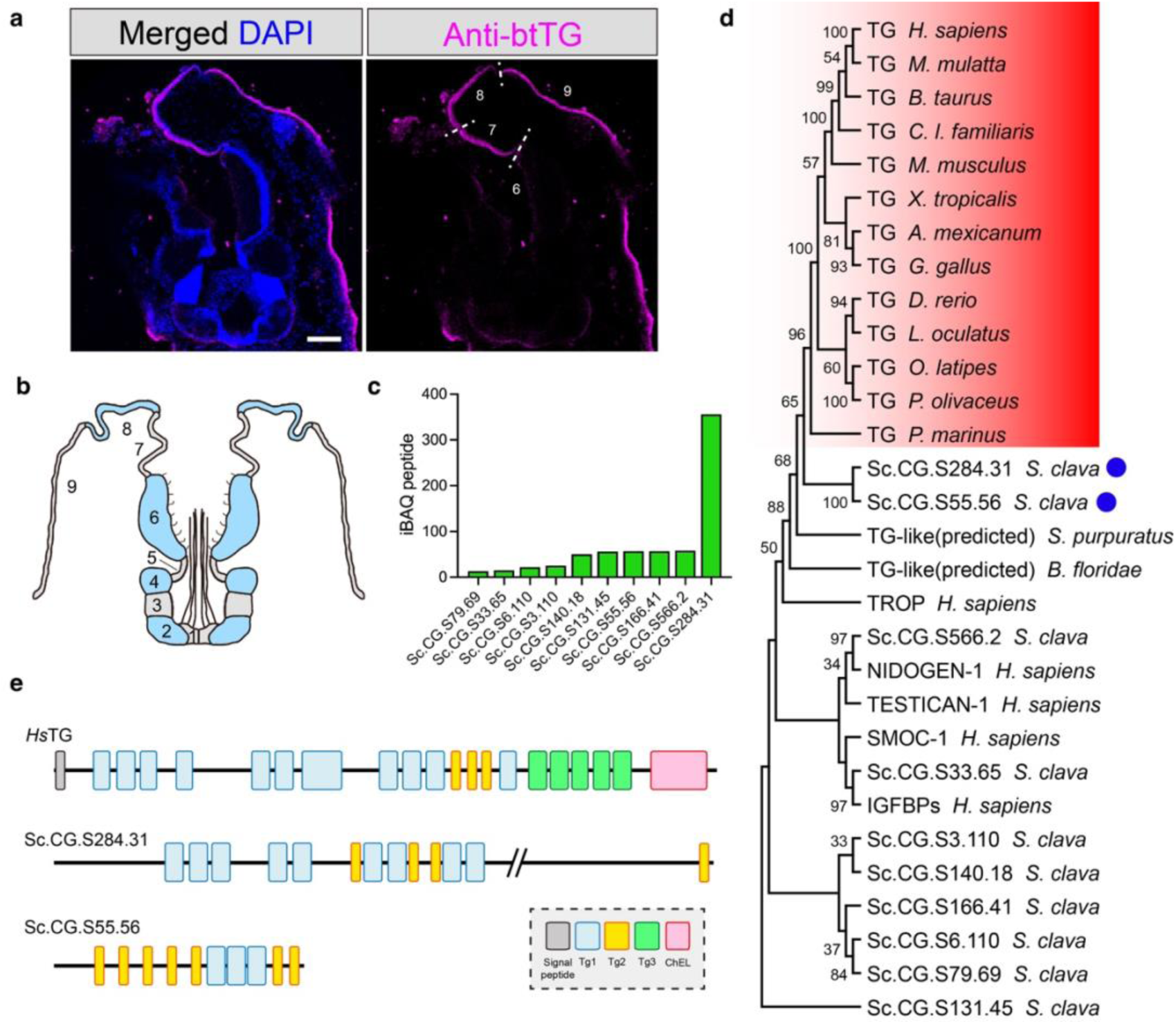
Identification of Thyroglobulin-like (TG-like) proteins in ascidian through immunoprecipitation and phylogenetic analysis. **(a)** Transverse section of endostyle of adult *S. clava* labeled by antibody of *Bos taurus* TG protein (magenta) and DAPI (blue). Scale bars = 100 μm. **(b)** Transverse section of endostyle of adult *S. clava*, refer to Hiruta et al^30^. The zones are labelled in grey (zone 1, 3, 5, 7 and 9) and blue (zone 2, 4, 6 and 8). **(c)** The iBAQ peptide number of Tg1-domain-containing proteins mapping to the dataset of immunolabeled proteins of Anti-*Bt*TG. Intensity represents the enrichment of the target protein among all immunolabeled proteins of Anti-*Bt*TG. **(d)** Phylogenetic analysis of TG proteins in vertebrates (in red box), the ascidian *S. clava* Tg1-domain-containing proteins in **(c),** the Tg1 domain-containing proteins in *Homo sapiens*, and the predicted TG-like protein in published literature ^9^. Bootstrap values greater than 30 are indicated in the nodes of the phylogenetic tree. The blue dots label the closest relative of vertebrate TG proteins among ascidian *S. clava* Tg1-domain-containing proteins in **(c). (e)** Domain distributions of human *Homo sapiens* TG (*Hs*TG), Sc.CG.S284.31 and Sc.CG.S55.56 proteins. Signal peptide is indicated in grey, Tg1 domains in blue, Tg2 in yellow, Tg3 in green, and ChEL in pink.

### Sc.CG.S284.31 is a TH protein precursor protein

We further investigated the expression patterns of genes encoding Sc.CG.S284.31 and Sc.CG.S55.56 proteins. Based on transcriptomic data from the embryonic and larval development of *S. clava*^27^, we revealed that *Sc.CG.S284.31* gene was up-regulated from tail-bud stage with the highest expression level during larval stages, which was similar to the temporal expression profiles of the homologs of vertebrate TH-synthesis-related genes, such as thyroid peroxidase (*ScTpo*), Sodium/iodide symporter (*ScNis*), thyroid transcription factors (*ScTtf1* & *ScTtf2*) (Fig. 2a). Distinct from the temporal expression profiles of *Sc.CG.S284.31* and *ScTpo*, *Sc.CG.S55.56* gene had the highest expression level at tail-regressed larval stage (Fig. 2a).

**Figure 2.**
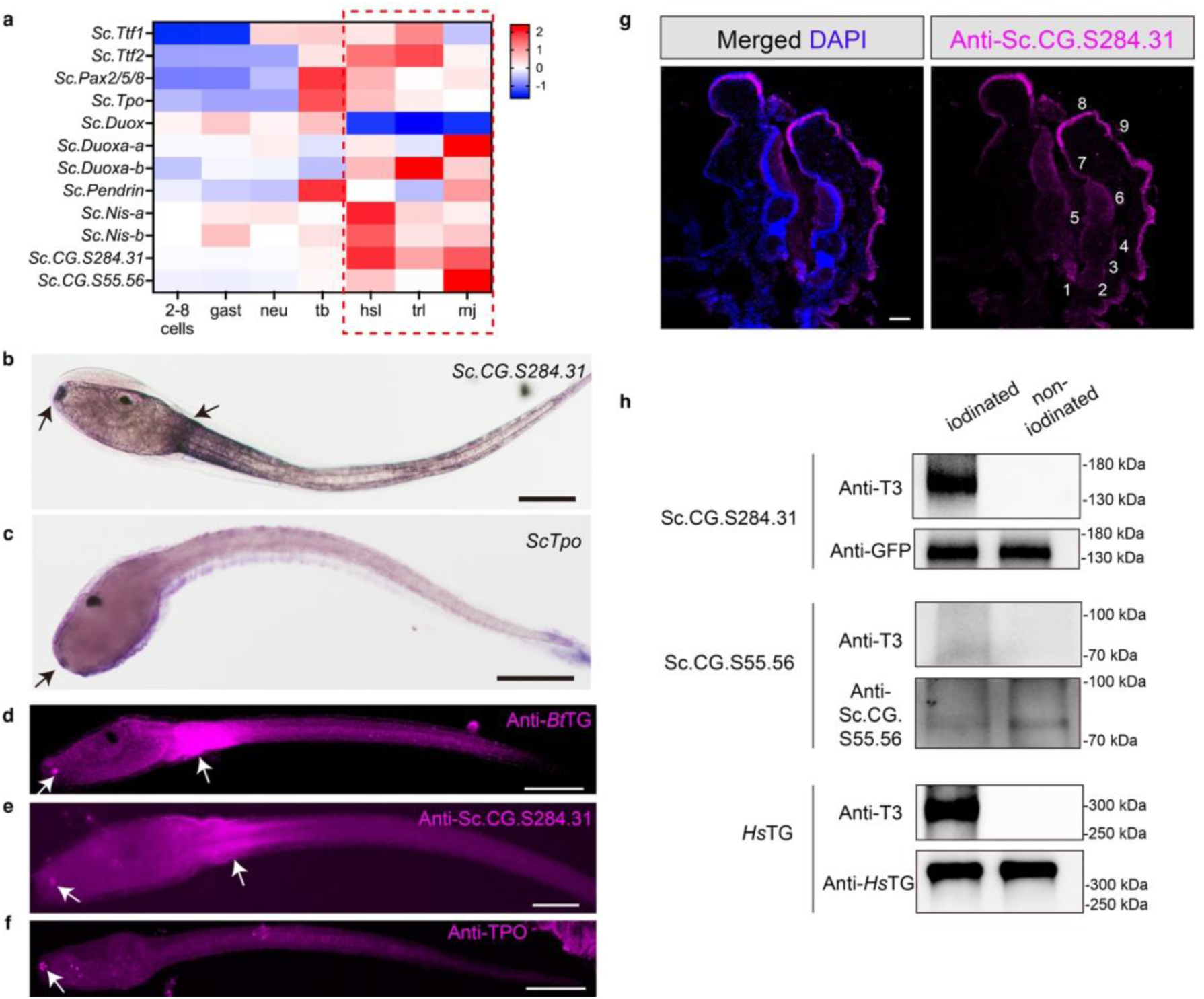
Expression patterns and *in vitro* functional verification. **(a)** Heat map shows the expression profiles of *Sc.CG.S284.31* and *Sc.CG.S55.56* genes, as well as the homologs of TH-synthesis related genes, during *S. clava* embryonic and larval development, including 2-8 cells (2-8 cells), gastrula (gast), neurula (neu), tailbud (tb), hatched swimming larval (hsl), tail regressing larval (trl), and metamorphic juvenile (mj) stage. The larval stages (including hsl and trl) and juvenile stage (including mj) are labeled in red box. **(b-c)** mRNA expression patterns of *Sc.CG.S284.31* (b) and *ScTpo* (c) in *S. clava* larvae, detected by *in situ* hybridization. The black arrows indicate the signals. Scale bars = 100 μm. **(d-f)** Immunofluorescence staining of *S. clava* larvae by antibodies of *B. taurus* TG protein (d), Sc.CG.S284.31 protein (e), and TPO protein (f), respectively. The white arrows indicate the signals. Scale bars = 100 μm. **(g)** Immunofluorescence staining of transverse section of adult *S. clava* endostyle by antibody of Sc.CG.S284.31 protein (magenta) and DAPI (blue). Scale bars = 100 μm. The transverse section of endostyle of adult *S. clava* is showed in Figure 1b, and the zones are indicated in the figure. **(h)** Validation of TH synthesis on the iodinated proteins by western blot. The segment of Sc.CG.S284.31 protein containing 9 Tg1 domains was used for *in vitro* iodination, with eGFP tag and a total molecular weight 160 kDa. The full length of Sc.CG.S55.56 protein was used for *in vitro* iodination, with 6×His tag and a total molecular weight 88 kDa. The *in vitro* iodination with human TG protein (330 kDa, with 6×His tag) are positive control. The *in vitro* iodination performing with PBS are negative controls.

The *in-situ* hybridization showed that the *Sc.CG.S284.31* transcripts were distributed in the anterior trunk and the epidermis between the trunk and tail (Fig. 2b, Supplementary Fig. 2a), exhibiting similar distribution pattern with *ScTpo* and THs (Fig. 2c, Supplementary Fig. 2b, Supplementary Fig. 3). In contrast, *Sc.CG.S55.56* transcripts were distributed in the posterior trunk (Supplementary Fig. 2c-d). These results therefore revealed that *Sc.CG.S284.31*have a similar expression dynamics and pattern as *ScTpo*, and might play a role in TH synthesis. We therefore prepared an antibody against this Sc.CG.S284.31 protein and performed immunofluorescence staining on the *S. clava* swimming larvae. The staining signals appeared at the anterior trunk and the epidermis between the trunk and tail, which was similar to distribution pattern of immunolabeled protein(s) through the anti-*Bt*TG staining (Fig. 2d-e). Furthermore, immunofluorescence staining with anti-TPO antibody revealed that ScTPO proteins were also present in the anterior trunk of *S. clava* larvae (Fig. 2f).

Given that these proteins were originally identified from the endostyle, which in ascidian adult is homologous to the thyroid follicles of vertebrates^24^, we investigated whether Sc.CG.S284.31 protein could be detected in the thyroid-equivalent region of *S. clava* endostyle. The immunostaining with antibody specific to Sc.CG.S284.31confirmed that this protein was specifically localized within zone 7 to 9 in endostyle (Fig. 2g), corresponding to the thyroid-equivalent region.

To determine whether Sc.CG.S284.31 can indeed serve as the matrix, providing hormonogenic tyrosine sites for TH synthesis, we performed immunoblotting assay to detect TH formation after *in vitro* protein iodination^10^ (Supplementary Fig. 4a). Given the extremely large molecular weight (630.45 kDa) of Sc.CG.S284.31 protein and the difficulty of cloning full-length sequence due to the presence of a large number of repetitive sequences, we constructed a vector to express recombinant Sc.CG.S284.31 protein containing Tg1-domain-enriched sequences (Supplementary Fig. 4b, Supplementary Table 1). The purified recombinant Sc.CG.S284.31 were used in iodination and subsequent immunoblotting assay. The results showed that T3 antibody readily detected specific bands between 130 to 180 kDa (for the recombinant Sc.CG.S284.31) or >300 kDa (for the recombinant human TG) after the proteins were iodinated (Fig. 2h). However, the T3 antibody failed to detect any signal in the iodinated sample of purified Sc.CG.S55.56 (Fig. 2h).

Overall, the results of expression analysis as well as the *in vitro* iodination experiment clearly demonstrated that Sc.CG.S284.31 protein, rather than Sc.CG.S55.56, could be a TH protein precursor providing hormonogenic sites for *de novo* TH production, similar to the function of vertebrate TG proteins. Therefore, we named Sc.CG.S284.31 as ScTG-like.

### *ScTg-like* is crucial for *in vivo* TH synthesis required for larval metamorphosis

To demonstrate whether *ScTg-like* is involved TH synthesis *in vivo*, we conducted RNA inference (RNAi) experiments to knockdown its expression and analyzed TH levels and larval metamorphosis (Fig. 3a). After treating larvae with specific siRNAs of *ScTg-like*, the expression of *ScTg-like* were significantly decreased in both mRNA and protein levels (Fig. 3b, Supplementary Fig. 5). We simultaneously measured the THs levels by ELISA. Both T4 and T3 levels in *S. clava* larvae were significantly decreased upon knockdown of *ScTg-like*, comparable to that of methimazole (MMI, a TPO inhibitor^28^)-treated larvae (Fig. 3c-d). Meanwhile, we observed that the tail regression of larvae was affected after knocking down *ScTg-like* (Fig. 3e-f), suggesting the crucial role of *ScTg-like* in tail regression, a typical event that occurs at the early metamorphosis of ascidian larva^27^. We further tracked the morphological changes of *S. clava* larvae with or without *ScTg-like* knock-down, and found that the process of metamorphosis of siRNA-treated larvae was largely delayed. In addition, compared with juveniles without *ScTg-like* knock-down, the “juveniles” in siRNA treated groups exhibited abnormal phenotypes with inextensible siphons and immature organs (Supplementary Fig. 6-7), suggesting that the organ morphogenesis of *S. clava* larvae during metamorphosis was regulated by *ScTg-like* gene. Interestingly, we demonstrated that the role of *ScTg-like* in metamorphosis is effectively linked to its role in TH synthesis, since, in rescue experiment, T3 treatment effectively restored the metamorphosis of *ScTg-like* knockdown group to a level comparable to those of the siRNA control and seawater control groups (Fig. 3g-i, Supplementary Fig. 6). Collectively, we established that the *ScTg-like* was indispensable for metamorphosis by functioning as the protein precursor for TH formation in *S. clava* larvae.

**Figure 3.**
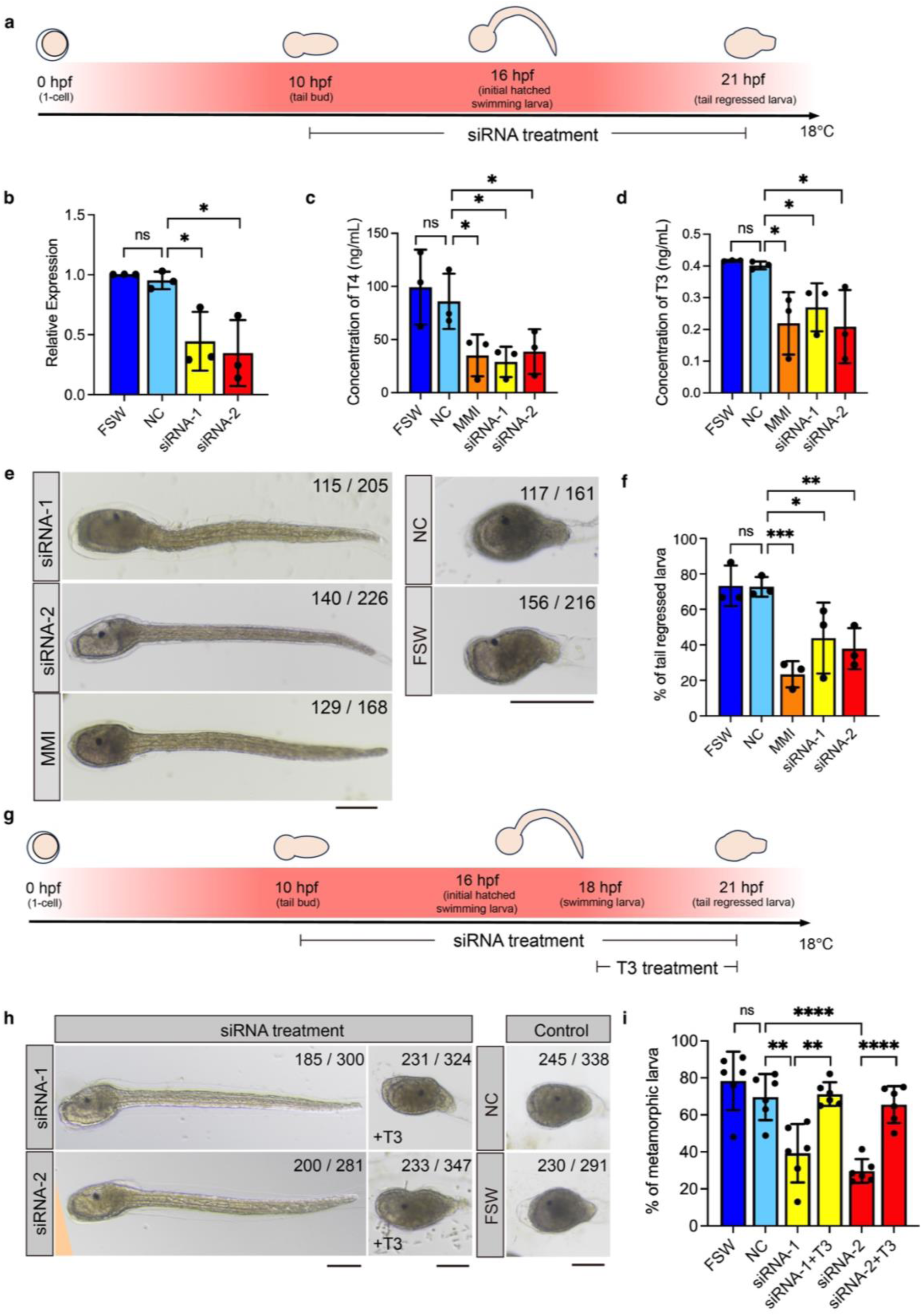
*In vivo* functional verification of *ScTg-like* in *S. clava* larvae. **(a)** Workflow of gene knockdown experiment in *S. clava* larva. The concentrations of siRNAs are 0.4 μM. **(b)** Quantifications of *ScTg-like* mRNA expression in gene knockdown (siRNA-1 and siRNA-2, two specific siRNAs of *ScTg-like* are designed to treat the *S. clava* larvae respectively), negative control siRNA treatment (NC, as the negative control), and filtered seawater treatment (FSW, as the blank control) groups through RT-qPCR method. *, *p*-value ≤ 0.05. ns, *p*-value > 0.05, no significant. **(c-d)** Quantification of T4 **(c)** and T3 **(d)** levels in gene knockdown (siRNA-1 and siRNA-2), MMI treatment (MMI, as the positive control, 120 mg/L), NC, and FSW groups. *, *p*-value ≤ 0.05. no significant (ns), *p*-value > 0.05. **(e)** Phenotypes of *S. clava* larvae in gene knockdown (siRNA-1 and siRNA-2), MMI, NC, and FSW groups. The fraction showed on each image indicates the number of *S. clava* with the phenotypes showed in the image / total number of *S. clava* in each experimental group. Scale bars = 50 μm. **(f)** Statistical results of tail regressed larvae in e. **, *p*-value ≤ 0.01. *, *p*-value ≤ 0.05. ns, *p*-value > 0.05. **(g)** Workflow of gene knockdown and rescue experiment in *S. clava* larva. The concentrations of siRNAs are 0.4 μM. The concentration of T3 for rescue is 50 μg/mL. **(h)** Phenotypes of *S. clava* larvae gene knockdown (siRNA-1 and siRNA-2), rescue (siRNA treatment + T3 rescue), NC, and FSW groups. The fraction showed on each image indicates the number of *S. clava* with the phenotypes showed in the image / total number of *S. clava* in each experimental group. Scale bars = 50 μm. **(i)** Statistical results of tail regressed larvae in h. ****, *p*-value ≤ 0.0001. **, *p*-value ≤ 0.01. ns, *p*-value > 0.05. All values in this figure are depicted as mean ± SEM. Significant tests are analyzed by T-test.

### A follicle-like structure functions as the thyroid-equivalent region for TH synthesis and storage in *S. clava* larva

In *S. clava* larvae, our anti-ScTG-like antibodies distinctly detected a signal in an invaginated structure consisting of four closely contacted cells, located at the central anterior region of the trunk (Fig. 2e and 4a-b). This invaginated structure could also be specifically labeled by the anti-TPO antibody (Fig. 2f and 4c), suggesting that the cells within this structure might function similarly to a vertebrate thyroid follicle. To validate this hypothesis, we conducted further immunostaining staining of THs (T3 and T4), ScDUOX1, and several transcriptional factors (including sc-FOXQ1 and sc-PAX2/5/8, whose vertebrate homologs are involved in thyroid morphogenesis and TH synthesis) in the larvae^29,30^. Notably, all the antibodies used in immunofluorescence experiment were able to distinctly label this invaginated follicle-like structure (Supplementary Fig. 8). Moreover, the peanut agglutinin (PNA) and wheat germ agglutinin (WGA) apparently labeled the follicle-like structure in *S. clava* larvae, indicating the glycoproteins were enriched in this structure (Supplementary Fig. 9a-c). Similar results could also be observed in the thyroid follicle of zebrafish (Supplementary Fig. 9d-f). Additionally, the anti-T3 signals were undetectable, as well as the anti-ScTG-like protein signals, in the follicle-like structure after knocking down the *ScTg-like* gene (Fig. 4d, Supplementary Fig. 8). The findings provide compelling evidence that this invaginated follicle-like structure, with expression of ScTG-like protein, THs, and other thyroid function-related proteins, serves as the site of TH synthesis and storage, representing a proto-thyroid-follicle in *S. clava* larvae.

**Figure 4.**
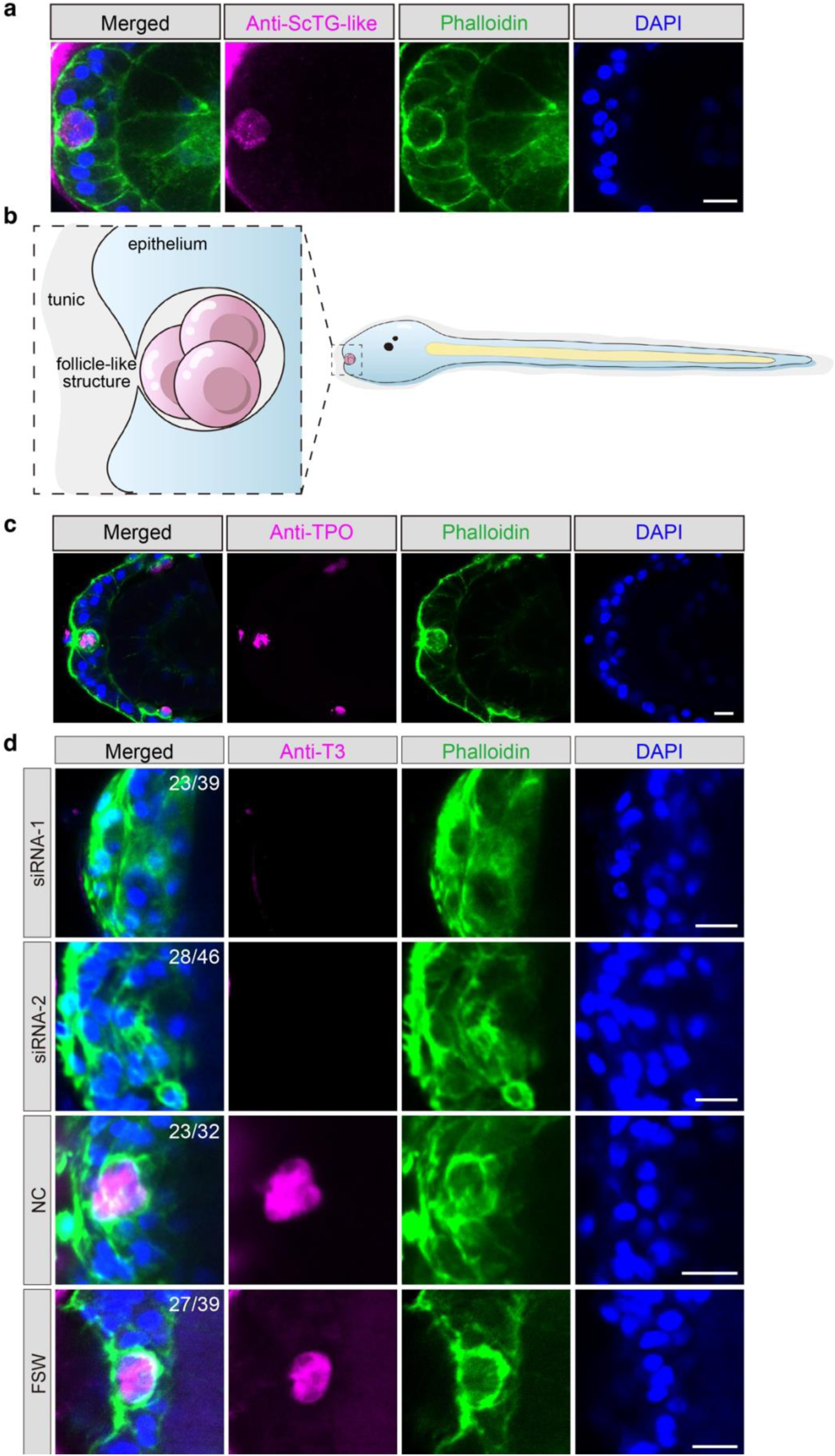
A follicle-like structure in *S. clava* larva functions in TH synthesis and storage. **(a)** The novel follicle-like structure, at the most anterior of *S. clava* larval trunk, is labeled by antibodies of ScTG-like protein (magenta), phalloidin (green), and DAPI (blue). Scale bars = 10 μm. **(b)** Diagram of the follicle-like structure at the central arterial of *S. clava* larval trunk. **(c)** The follicle-like structure is labeled by antibody of TPO protein (magenta), phalloidin (green), and DAPI (blue). Scale bars = 10 μm. **(d)** The abolishment of TH synthesis inside of the follicle-like structure after knocking down of *ScTg-like* gene. Two specific siRNAs, siRNA-1 and siRNA-2, was used to knock down the expression of *ScTg-like* gene respectively. Negative control, larvae treated with NC siRNAs. Blank control, larvae treated with FSW. The concentrations of siRNAs are 0.4 μM. The presence of T3 signals in the follicle-like structures was detected using antibody of T3 (magenta). The phenotypes of the follicle-like structures were labeled by phalloidin (green) staining. The nuclei are labeled using DAPI (blue). Scale bars = 5 μm. The fraction showed on each image indicates the number of *S. clava* with the phenotypes showed in the image / total number of *S. clava* in each experimental group.

### Structural conservation suggests functional homologues across bilaterians

Despite extensive homology searches, we did not identify a clear orthologue of *ScTg-like* in any other species, including other tunicates such as the closely related *Styela plicata*. This likely reflects the rapid evolution of tunicate genomes^31^ and the low primary sequence conservation typical of TG proteins. Indeed, as previously observed in vertebrates^4^, the conservation of TG function appears to rely more on structural architecture than on sequence similarity. ScTG-like protein shares hallmark features with vertebrate TGs, including large size (>2000 aa), multiple Tg1 domains, extensive disulfide bridges, and a C-terminal region devoid of disulfides that may fulfill a ChEL-like role. Using this domain-based signature, we identified several candidate proteins that match the predicted structural organization of ScTG-like protein, not only in other tunicates but also in cephalochordates, echinoderms, hemichordates and protostomes (Supplementary Fig. 10). Although precise sequence-based comparisons remain difficult, these findings support the existence of functional homologues of TG-like proteins across bilaterians—in accordance with the notion that endogenous TH synthesis may be an ancient and conserved feature of this clade.

Taken together, our results demonstrated the existence in *S. clava*, an urochordate, of a protein encoded by *ScTg-like* gene that play a key role as a thyroid hormonogenic protein localized in the follicle-like structure of ascidian tadpole larva. These data provide novel insights into the molecular mechanisms of TH synthesis outside vertebrates and contributes to understanding the evolutionary trajectory of the TG protein.

## Discussion

THs are crucial factors for metamorphosis of both vertebrates and invertebrates^4,15^. The mechanism of TH synthesis has been well documented in vertebrates, but remains elusive in invertebrates. This is due to the fact that TG homologs or functional alternatives in invertebrate have not been exactly identified despite that many orthologues of genes involved in TH synthesis are already present in these species^4,6,30,32^. In 1978, based on the positive immunofluorescence results with Anti-*Bt*TG in endostyle of Conklin’s ascidian *S. clava* that was first used by Conklin for embryonic pattern organization and cell lineage tracking^33^, it has been proposed that the vertebrate TG protein homolog might exist in *S. clava* endostyle^25^. However, over the past decades, no definitive conclusion has been made regarding whether invertebrates really process the TH precursors (e.g hormonogenic matrix proteins) that directly participate in TH synthesis, as found in vertebrates. Here, we identified a novel high molecular weight protein ScTG-like in *S. clava*, which only processes Tg1 domains but exhibit similar function as vertebrate TG proteins. Both *in vitro* and *in vivo* studies demonstrated that ScTG-like is a protein precursor that directly participates in TH synthesis as a hormonogenic matrix protein in ascidian. Furthermore, based on the expression of ScTG-like and other TH synthesis-related proteins, we discovered a follicle-like structure in the anterior of larva trunk, functioning as a proto “thyroid gland”. By identifying and characterizing a molecule that serves as the protein precursor for endogenous TH synthesis in a non-vertebrate chordate, our work provides important new insights into the mechanisms of hormone production in these organisms.

Iodine is a key element for THs synthesis, and it exhibits alive metabolic activity in marine organisms, affecting their life processes^18,34,35^. For example, marine bacteria, kelp and sea urchin uptake iodine from seawater through the process of hydrogen peroxide-dependent diffusion^36–38^. Although the presence of THs and their derivatives, as well as iodoproteins, in marine organisms have been demonstrated by several previous biochemical studies^19,34,39,40^, it has not yet conclusively established that the *de novo* TH synthesis on a matrix protein effectively occurs in these species. Indeed two alternative possibilities cannot be reasonably ruled out: 1) these species may utilize intermediates sourced from environment or symbiotic organisms to generate THs without the involvement of endogenous hormonogenic matrix protein^16,41,42^, and 2) the previously detected iodoproteins in endostyle extracts may be TH-binding proteins or TH-transporter proteins for hormone transport (e.g. thyroxine-binding globulin and many types of membrane transporters^43^), rather than proteins directly involved in TH production. With the convincing evidence that ScTG-like protein is directly involved in TH synthesis within the follicle-like structure of larvae and specific zones of adult endostyle (Fig. 2g-h and 4a), we are confident to conclude that the process of TH biosynthesis in invertebrates, at least for ascidian *S. clava*, is generally analogous to that in vertebrates, using a protein similar but not directly related to vertebrate TG. This clearly suggests that the *de novo* TH synthesis process is an ancestral and conserved pathway.

The identification of TG-like proteins with conserved domain architectures in non-vertebrate chordates—resembling the vertebrate TG layout but lacking the ChEL domain—raises the possibility that key structural elements of TH precursor proteins may have deeper evolutionary origins, potentially extending to other bilaterian lineages (Supplementary Fig. 10). While this does not constitute definitive evidence, it is consistent with the hypothesis that TG-like proteins, and possibly endogenous TH synthesis, could represent an ancient bilaterian innovation. These findings suggest that other metazoan phyla, such as annelids or molluscs, may harbor structurally conserved but sequence-divergent TG-like proteins, offering new avenues to investigate the synthesis, regulation, and function of TH in invertebrates. Although large-scale synteny is unlikely to be preserved in tunicates, due to their highly rearranged genomes^31^, examining genomic context and expression patterns—particularly during metamorphosis—in other bilaterians could provide supporting evidence. In addition, structural modeling approaches (e.g., AlphaFold) may help uncover conserved folding and function despite low similarities of primary sequences, contributing to our understanding of the evolutionary emergence of TH signaling and its potential role in the diversification of metazoan life cycles. Additionally, proteins containing the ChEL domain have also been identified in invertebrate deuterostomes, and studies have shown that these genes undergo gene duplication and rearrangement, resulting in proteins with diverse functions^44^. Investigating the interactions between the ChEL domain-containing proteins and TG-like proteins, as well as the potential for gene fusion, may provide insights into the evolutionary transition from invertebrate *Tg-like* genes to vertebrate *Tg* gene.

Thyroid follicles serve as the functional units in TH production and storage^45^. In vertebrates, thyrocytes surround a single layer of epithelial cells, with the lumen filled with colloid at the center^45^. Secreted proteins are synthesized in follicular cells and transported into the colloid. Interestingly, the follicle-like structure identified in *S. clava* larva exhibited morphological and functional similarities to thyroid follicles (Fig. 4b). The follicle-like structure was observed at the most anterior of *S. clava* larval trunk, surrounded by epithelial cells (Fig. 4b). It was opened outward to connect with the outer tunic compartment (Fig. 4b). Glycosylation in the vertebrate thyroid gland is thought to be essential for its function^6,46^. High glycoprotein content reflects the high secretory activity of a tissue or structure (e.g. follicle structure) since most secretory proteins need to be glycosylated before entering the secretory pathway. Vertebrate TG proteins are the typical glycoproteins, and their glycosylation is essential for secretion and function^5,6^. Thus, staining for glycoproteins can be used to reflect the function of thyroid gland^47^. Thus, we found that both the follicle-like structure in *S. clava* larva and the thyroid follicle in zebrafish were apparently labeled by PNA and WGA, indicating that both of these two structures are enriched with glycoproteins (Supplementary Fig. 9). Moreover, knockdown of *ScTg-like* completely eliminated TH production within follicle-like structure (Fig. 4d), further indicating the functional similarity between the thyroid follicles and *S. clava* follicle-like structure. Furthermore, the genes involved in TH synthesis were expressed both in the follicle-like structure in larvae and thyroid equivalent region of endostyle in adult (Supplementary Fig. 8 and 11). In addition, a similar structure was also detected in the larvae of ascidian *Herdmania curvata*, which is filled with the Hemps (a key protein that regulates metamorphosis), indicating the crucial function of the follicle-like structure in metamorphosis of ascidians^48^.

Previous studies have proved that TH play crucial roles in tail regression of ascidians^49^. In this study, we found that after reducing the TH levels, not only the tail regression, but also the subsequent metamorphic process was impacted (Figure 3, Supplementary Fig. 6). After knocking down the expression of *ScTg-like* gene, the process of metamorphosis was significantly delayed, and the morphology of “juvenile” was abnormal, with the inextensible siphons and immature organs (Supplementary Fig. 6). Considering that the *ScTg-like* gene contains a length of 17,373 base pairs coding sequence (Supplementary data 1), it is difficult to clone its full-length coding sequence and synthesize mRNA for rescue experiment. Instead, we performed the rescue experiment by T3 treatment, which clearly showed that the rescued larvae developed normally, similar to the wild type (Figure 3, Supplementary Fig. 6). These results suggest that TH signal is of great importance in controlling the timing of metamorphosis, morphogenesis, and organ formation in *S. clava* larvae. In vertebrates, TH exerts its regulatory functions by binding to thyroid hormone receptors (TRs), which activates downstream genes^2^. However, whether a similar mechanism exists in invertebrates remains unknown. In amphioxus, T3 derivative TRIAC is the ligand of TR, which supports that TRIAC is the active form of the hormone ^50^. In *S. clava*, *Tr* mRNA is mainly expressed at the anterior end of the trunk and in the tail epidermis of larvae^27^, which appears to coincide with the distribution of T3 (Supplementary Fig. 3). However, it remains to be determined whether these two molecules can actually bind.

Altogether, our study provided the direct evidence of endogenous *de novo* TH synthesis outside vertebrate through identifying the TG-like protein in proto-vertebrate ascidian, which solved a long-standing mystery about the evolutionary “missing” TH protein precursor outside vertebrates. Our findings also shed lights on the evolutionary trajectory of the TG protein, and the emergence of a major endocrine system in vertebrates.

## Methods

### Sample collection

#### Cyrosections

Endostyle tissue was isolated from *S. clava* adult. The head was isolated from adult zebrafish (*Denio rerio*). The samples were subsequently fixed in 4% paraformaldehyde (PFA) at room temperature (RT) for 2 hours. Following three washes with phosphate-buffered saline (PBS), the tissue was treated with 30% sucrose solution in PBS. The sedimented endostyle was then embedded in Tissue-Tek OCT compound (Sakura, Torrance, CA, 4583) and snap-frozen in liquid nitrogen. The sample was stored at -80°C before cryosection. Prior to sectioning, the OCT-embedded endostyle was equilibrated by placing it in a -20°C freezing microtome for 30 minutes. Cryosections were collected at 10 μm intervals along the transverse plane utilizing a cryostat (CM3050, Leica). Subsequently, the sections were mounted onto adhesive slides.

#### Larvae

The fertilization and culture of *S. clava* embryos were performed according to the previously detailed procedures by Wei et al.^27^ and Lin et al.^51^. The larvae of *S. clava* were collected and fixed by 4% PFA at RT for 2 hours. Following washing with PBS for three times to eliminate PFA, the larvae were stored in 4°C.

### Staining

#### Immunofluorescence

For immunofluorescence staining, the samples were initially permeabilized through washing with PBST (0.1% Triton X-100 in PBS) for five times within 8 hours. Subsequently, blocking was performed using 10% goat serum at RT for 1 hour. The samples were then incubated with primary antibodies (1:100 to 1:50) overnight at 4°C with shaking. Following this, the samples were washed by PBST for five times within 8 hours. Subsequently, samples were incubated with secondary antibody (A21422, Invitrogen) and Phalloidin 488 (A12379, Invitrogen) overnight at 4°C with shaking. After repeating the washing step with PBST under the same conditions, the samples were mounted with mounting solution (Vector Laboratories) containing DAPI and observed by a confocal microscope (Zeiss, LSM900). The primary antibodies used in this study were listed in Supplementary table 2.

#### Peanut agglutinin (PNA) staining

The samples were initially permeabilized through washing with PBST for five times within 8 hours. Subsequently, blocking was performed using 10% goat serum at RT for 1 hour. The samples were then incubated with biotinylated PNA (1:300) overnight at 4°C with shaking. Following this, the samples were washed by PBST for five times within 8 hours. Subsequently, samples were incubated with Vari Fluor 488-Streptavidin (HY-D1808, MedChemExpress) and TRITC Phalloidin (40734ES75, Yeasen) for two hours at RT with shaking. Following this, the samples were washed by PBST for three times within one hour. Subsequently, the samples were mounted with mounting solution (Vector Laboratories) containing DAPI and observed by a confocal microscope (Zeiss, LSM900).

#### Wheat germ agglutinin (WGA) staining

The samples were initially permeabilized through washing with PBST for five times within 16 hours. Subsequently, samples were incubated with WGA (1:50) for three days at RT with shaking. Following this, the samples were washed by PBST for five times within 8 hours. Subsequently, the samples were mounted with mounting solution (Vector Laboratories) containing DAPI and observed by a confocal microscope (Zeiss, LSM900).

### Immunoprecipitation coupled with mass spectrometry (IP-MS)

The endostyle tissues were isolated from *S. clava* adults, and subsequently shredded and homogenized. The homogenized tissues were incubated with RIPA lysis buffer, supplemented with 1% PMSF to effectively inhibit protease activity, for 20 mins on ice. Following this, high-speed centrifugation was carried out at 12,000 rpm for 5 mins at 4°C. The supernatant was then collected, and the protein concentration was determined using the Bicinchoninic Acid (BCA) Assay. Subsequently, the endostyle protein was incubated with the antibody specific to *B. taurus* TG protein (Supplementary table 2) overnight at 4°C with shaking, to facilitate the formation of the antigen-antibody complex. Magnetic beads were then introduced into the system and incubated for 2 hours at room temperature, allowing the formation of the antigen-antibody-beads complex. Magnetic adsorption was subsequently performed using a magnetic rack to enrich the antigen-antibody-beads complex. After removing the supernatant, the complex was washed three times with PBS to ensure purity. Ultimately, the complex was collected for subsequent mass spectrometry identification.

### Bioinformatics analysis

#### Protein domain prediction for S. clava genome

HMMER was employed with profile Hidden Markov Models (pHMMs) from the Pfam database, which contains domain-specific pHMMs representing conserved protein families, to predict the domain distribution of protein sequences in the whole genome of *S. clava*^27^. The hmmscan module was executed using default parameters (e-value ≤ 1e-5) to scan each protein sequence against the pHMM library, prioritizing domain coverage and statistical significance. The Proteins containing Tg1 domain(s) (Pfam ID: PF00086.23) were identified according to the results of protein domain prediction. Predicted domains were manually validated through cross-referencing with SMART database annotations (http://smart.embl-heidelberg.de).

#### Phylogenetic analysis

The information of protein sequences for phylogenetic analysis was showed in supplementary files (Supplementary table 3). Sequences were aligned with MEGA software (version 11.0) using the ClustalW with default parameters followed by manual refinement to optimize conserved motif alignment. The final alignment was used for phylogenetic analysis employing the Neighbor-Joining (NJ) method under the *p*-distance model. Branch support was assessed via 1000 bootstrap replicates, with nodes considered significant at > 30% bootstrap values.

#### Downloading and preparing genomes

DNA assemblies were obtained via NCBIDatasets CLI v18.3.1. For each organism selected from the taxa Tunicata (txid 7712), Cephalochordata (txid 7735), Petromyzontiformes (txid 7745), Annelida (txid 6340), Echinodermata (txid 7586), Hemichordata (txid 10219) and Mollusca (txid 6447), the most complete assembly level available was downloaded.

#### Generation or collection of proteomes

When the proteome was available, the *.faa file provided by RefSeq/GenBank was used as is. If absent, GeneMark ES v4.73 predicted ORFs *de novo*.

#### Reference vertebrate sequences and construction of HMM profiles

Thirty vertebrate sequences (25 RefSeqs in the taxa Amphibia (txid 8292), birds (txid 8782), bony fishes (txid 7898), Squamata (txid 8509), Testudines (txid 8459), and 5 mammalia (txid 40674) UniProtKB/Swiss Prot: THYG_HUMAN, THYG_MOUSE, THYG_RAT, THYG_PIG, THYG_BOVIN) were used as a reference set. The 5 annotated Uniprot sequences were used to create a first version of the HMM profile for the Tg1, Tg2, Tg3, and ChEL domains. Then, using hmmsearch (default threshold), the domains were identified in the other 25 sequences. All domain sequences were extracted and aligned with MAFFT v7.525 (--localpair --maxiterate 1000) and then converted into specific HMM profiles via hmmbuild (HMMER v3.4). These are the profiles that will be used for the rest of the pipeline.

#### Annotation of invertebrate proteomes

For the detection of relevant proteins, three methods were combined: (1) HMM profile search, using hmmsearch (default threshold) for the detection of Tg1, Tg2, Tg3 and ChEL domains. Only sequences with at least 5 Tg1 motifs were retained by this method. (2) Search by regex, the search patterns are Tg2-like domain (C××C×G) and Tg2-like domain (C××C××G), potential disulfide bridges C (1-20) C, and tyrosine residues Y. Blastp v2.16.0+ search against each taxon with lamprey TG protein and *S. Clava* TG-like protein as reference sequences. HMM profile search and regex search methods were applied to the blastp results to create an annotation file. Each domtbl annotation file was converted to gff3 format.

#### Visualization of protein architecture

To graphically represent the architecture of candidate TG-like proteins, a Python script was developed using the Biopython (for sequence manipulation), Pandas (metadata management), and Matplotlib (visualization) libraries. It takes as input a fasta file with all the sequences (Supplementary table 4), a folder containing all the annotations in gff3 format, and a tsv file listing the metadata for each sequence. For each sequence, it generates three parallel plots: positions of the Tg/ChEL domains, potential disulfide bonds, and tyrosine residues, and generates a svg vector file and a png file for each sequence.

### Polymerase Chain Reaction (PCR) and RT-qPCR

Total RNA was extracted from the fresh samples utilizing the TRIzol method and subsequently dissolved in RNase-free ddH_2_O. cDNA synthesis was performed through reverse transcription employing PrimeScript II RTase (6210A, Takara). The newly synthesized cDNA served as the template for PCR analysis, employing specific primers of targeting genes (Supplementary table 5). To ensure accurate normalization, the gene expression level of *Sc18s rRNA* was utilized as an internal reference. PCR reactions were performed under the following conditions: pre-denaturation (3 min at 95°C) for one cycle, denaturation (15 s at 95°C)-annealing (15 s at 60°C)-extension (60 s/kb at 72°C) for 35 cycles, and complete extension (5 mins at 72°C) for one cycle. PCR products were analyzed by 1 % agarose gel electrophoresis. RT-PCR reactions were performed under the following conditions: pre-denaturation (3 min at 95°C) for one cycle, cyclic reaction (10 s at 95°C and 30 s at 58°C) for 40 cycles, melting curve (15 s at 95°C, 60 s at 60°C and 15 s at 95°C) for 1 cycle. The RT-PCR experiment was performed using Roche LC96. Data analysis was performed through 2^-ΔΔCt^ method. The primers for these experiments were listed in Supplementary table 5.

### *In situ* hybridization

Specific segments of the target genes were selected for the synthesis of probes. The primers utilized for amplifying specific segments were listed in Supplementary Table 5. The transcriptase binding sequence (SP6 RNA polymerase priming: 5’ ATTTAGGTGACACTATA 3’; T7 RNA polymerase priming: 5’ TAATACGACTCACTATAGGG 3’) was added to the 5’ end of the primer. Digoxigenin (DIG)-RNA probes were generated through *in vitro* transcription using either SP6 RNA polymerase (EP0131, Thermo Fisher Scientific) or T7 RNA polymerase (EP0111, Thermo Fisher Scientific), and subsequently purified using the LiCl and 100% ethanol. These DIG-RNA probes were stored at -80°C prior to hybridization.

The *in situ* hybridization procedure was conducted as previously described by Yasuo et al^52^. Briefly, samples were digested with proteinase K (15 μg/mL) for 30 mins at 37°C, followed by washing with PBT (0.1% Tween-20 in PBS). Samples were then refixed with 4% PFA (in PBT) for 30 mins at RT and washed again with PBT. Pre-hybridization was carried out using pre-hybridization buffer (50% formamide, 5× saline sodium citrate buffer (SSC), 0.1% Tween 20) for 15 mins at RT, followed by hybridization buffer (50% formamide, 5× SSC, 0.1% Tween 20, 5× Denhardt’s solution, yeast tRNA (100 μg/mL), and heparin (50 μg/mL)) for 3 hours at 50°C. Hybridization was subsequently performed with 2 ng/μL probes in hybridization buffer overnight at 50°C. After hybridization, samples were washed four times with wash buffer I (50% formamide, 5× SSC, 0.1% Tween 20), four times with wash buffer II (50% formamide, 0.5× SSC, 0.1% Tween 20), and four times with PBT. Samples were then blocked using 1% blocking buffer for 1 hour at RT. Following this, samples were incubated with DIG antibody overnight at 4°C. Subsequently, samples were washed four times with PBT and twice with TMNT (0.1 M Tris-HCl (pH = 9.5), 0.1 M NaCl, 0.02 M MgCl2, 0.1% Tween 20). Chromogenic detection was achieved using a Nitro blue tetrazolium (NBT)/bromochloroindolyl phosphate (BCIP) solution (1:50 diluted with TMNT).

### Protein expression and purification

To improve the expression efficiency in mammalian cells, we codon-optimized the sequence of *Sc.CG.S284.31* gene (Supplementary Data 1). The Tg1-enriched fragments of the *Sc.CG.S284.31* gene were cloned from the codon-optimized sequences, and subsequently subcloned into a linearized pEGFP-N1 vector (pre-linearization using *NheI* and *KpnI* restriction enzymes (Thermo Fisher Scientific)). Additionally, the full-length human TG-encoding sequence was cloned from a commercial plasmid obtained from the DNASU Plasmid Repository (Clone ID: HsCD00867406) and subcloned into a linearized pcDNA3.1(+) vector (pre-linearization using *NotI* restriction enzymes (Thermo Fisher Scientific)). The full-length Sc.CG.S55.56 protein-encoding sequence was cloned from cDNA library of *S. clava* and subcloned into a linearized pcDNA3.1(+) vector (pre-linearization using *NotI* restriction enzymes (Thermo Fisher Scientific)). The primers for clone were listed in Supplementary table 5. Purification of the resulting fragments was performed using the GeneJEL Kit (K0691, Thermo Fisher Scientific), ensuring high-quality material for subsequent experiments. Homologous recombination was carried out using the in-fusion method (C115, Vazyme). The constructed plasmids were transformed into DH5α competent cells. Bacteria successfully transformed with constructed plasmids were inoculated to LB media. Plasmid extraction and purification was performed using EndoFree Mini Plasmid Kit (DP123, Tiangen biotech).

HEK293T cells were employed for the *in vitro* expression of the recombinant protein. These cells were cultured in DMEM, supplemented with 10% fetal bovine serum, at 37°C in a 5% CO_2_ incubator. Twenty-four hours prior to transfection, the cells were plated in 10 cm cell culture dishes. Transient transfection was performed using Lipofectamine 3000 transfection reagent (L3000001, Invitrogen), following the manufacturer’s protocol. Twenty-four hours after transfection, cells were washed with PBS and cultured with serum-free DMEM for an additional twenty-four hours. Ultrasonication and ultracentrifugation techniques were employed to collect the recombinant Sc.CG.S284.31 protein and Sc.CG.S55.56 protein from cells. For the human TG protein, the secreted protein present in the medium of transfected cells was harvested for subsequent purification steps.

Protein purification was performed using Anti-GFP affinity gel (HY-K0229, MedChemExpress) for recombinant Sc.CG.S284.31 protein, and Ni Sepharose 6 FF (17531801, Cytiva) for Sc.CG.S55.56 protein and human TG protein. Prior to incubation, the agarose beads were thoroughly washed with PBS three times. The protein samples were then incubated with the agarose beads in gravity chromatography columns at 4°C overnight. The protein solvent was flowed out at a controlled rate of 1 mL/min. Subsequently, the agarose beads were washed three times with wash buffer (50 mM NaH_2_PO_4_, 300 mM NaCl, 20 mM imidazole, final pH = 8.0, filter with 0.22 μm membrane). The recombinant proteins were then eluted using elution buffer (50 mM, NaH_2_PO_4_, 300 mM NaCl, 500 mM imidazole, final pH = 8.0, filter with 0.22 μm membrane). To exchange the protein from the elution buffer to PBS, Sephadex DeSalting Gravity Columns (C500090, Sango) were employed, following the manufacturer’s protocol. Finally, the concentration of the protein sample was determined using the BCA method.

### *In vitro* enzymatic iodination and western blotting

The *in vitro* iodination experiment was performed following the protocol previously outlined by *Citterio* et al.^53^. The iodination mixture comprised 30 ng/μL lactoperoxidase, 2 μg/μL glucose, 0.352 ng/μL glucose oxidase, 100 μM NaI, and 150-200 ng/μL of purified protein. All reagents were purchased from Sigma. Subsequently, the sample was incubated at 37°C for 1.5 hours, and enzymatic reaction was stopped by addition of gel sample buffer and boiling for 10 mins.

Western blotting experiment was performed to detect the TH signal of the iodinated proteins. The iodinated samples were subjected to reducing SDS-PAGE, and transferred to 0.45 μm PVDF membrane (Millipore, MA, USA). Blocking was performed using 5% fat-free milk at room temperature (RT) for 1 hour. Subsequently, the membranes were incubated overnight at 4°C with a 1:1000 dilution of T3 Monoclonal Antibody (3A6) (MA1-21669, Invitrogen). Following washes with TBST (TBS containing 0.05% Tween-20), the membranes were incubated with species-specific HRP-conjugated secondary antibodies diluted at 1:2000 in blocking buffer at RT for 2 hours. After additional washes with TBST, the specific bands were visualized using a chemiluminescence digital imaging system (ImageQuant Las-4000 Mini, GE Healthcare). To detect the signals of proteins in iodinated samples, the anti-GFP rabbit polyclonal antibody (T0006, Affinity), anti-hTG mouse polyclonal antibody (A00186, GenScript) and anti-Sc.CG.S55.56 mouse polyclonal antibody (homemade) were employed (Supplementary table 2).

### Preparation of anti-Sc.CG.S55.56

The antibody of Sc.CG.S55.56 protein was made in our lab. The full-length Sc.CG.S55.56 protein -encoding sequence was cloned from cDNA library of *S. clava* (Supplementary table 5), and subcloned into a linearized pET30a vector (pre-linearization using *BamHI* and *EcoRI* restriction enzymes (Thermo Fisher Scientific)). Purification of the resulting fragments was performed using the GeneJEL Kit (K0691, Thermo Fisher Scientific), and homologous recombination was carried out using the in-fusion method (C115, Vazyme).

To obtain the Sc.CG.55.56 protein, prokaryotic expression using bacteria was performed. The recombined plasmids were transformed into BL21(DE3) competent cells. Bacteria successfully transformed with recombined plasmids were inoculated to LB solid media. Monoclonal colony was used for amplification culture in LB liquid media at 37°C with 220 rpm. When the OD value reached to 0.6, IPTG was added into bacterial fluid with the final concentration of 0.5 mM. The bacterial fluid was then developed in 20°C environment at speed of 160 rpm for 16 hours. Ultrasonication was used for extraction of total proteins in bacteria. The coomassie blue staining was performed to labeled the Sc.CG.S55.56 protein. The gels containing Sc.CG.S55.56 protein were split, and were homogenate through ultrasonication. To obtain the antibodies of Sc.CG.S55.56, intraperitoneal injection in mice was carried out once a week for a total of four times. A total of 200 μg protein was used for intraperitoneal injection every time. Finally, the blood of mice was drawn and further separated to obtain serum. The serum contains antibodies and is used in subsequent experiments.

### RNA inference and rescue experiment

Fertilized eggs were cultured at 18°C until the early tail bud stage, the embryos were gathered and placed into the wells of a 24-well plate, which had been pre-coated with 0.1% BSA. Each well contained more than 50 embryos. The siRNAs were designed and synthesized by Genecefe Biotechnology Co., Ltd (Wuxi, China). The sequences of siRNAs for this experiment were listed in Supplementary table 6. The sequence of negative control siRNA does not target any gene in the *S. clava* genome. The siRNAs were dissolved in RNase-free water to achieve a stock concentration of 20 μM. For transfection, the riboFECTTM CP transfection kit (C10511-05, Ribo Bio) was used according to the manufacturer’s instructions. The working concentration of the siRNAs for transfection was set at 0.4 μM, and the embryos were incubated with the siRNAs at 18°C for eleven hours. For rescue experiment, the T3 was added at 18 hpf with working concentration at 50 μg/mL. Following the incubation period, the phenotypes of the larvae were recorded using a microscope, and the percentages of metamorphic larvae were analyzed. Subsequently, the larvae were collected for RNA extraction.

### Enzyme linked immunosorbent assay (ELISA)

The larvae were dissociated using ultrasonication, and the ultracentrifugation was performed to obtain the supernatant for ELISA detection. Standardize each group of samples by weighing. The 25 μL of sample was incubated with 100 μL TH-HRP conjugate in each well of 96-well plate for 1 hour at RT. After incubation, the content of the wells was discarded, and each well was washed with 300 μL of wash buffer for five times. Subsequently, 100 μL of TMB substrate solution was added to each well, followed by incubation at RT in the dark for 15 mins. Then, 100 μL of stop solution was added to each well, and the plate was gently shaken. The absorbance of the samples was measured at a wavelength of 450 nm.

### Statistical analysis

All values are depicted as mean ± SEM. Significant test was analyzed by T-test. Data were estimated to be statistically significant at *p*-value ≤ 0.05. ****, *p*-value ≤ 0.0001. ***, *p*-value ≤ 0.001, **, *p*-value ≤ 0.01. *, *p*-value ≤ 0.05.

## Data Availability

The datasets presented in this study can be found in publicly available repositories. The transcriptomic data of *S. clava* during early developmental processes are deposited in NCBI SRA database (accession number PRJNA1232216). The Stereo-seq datasets on the *S. clava* endostyle have been deposited at China National GenBank DataBase (CNGBdb, https://db.cngb.org/) under the accession number CNP0004228.

## Acknowledgements

We sincerely thank Prof. Michael C. Thorndyke from Queen Mary University of London, Prof. Guoqing Wang and Dr. Xuan Zhu in Shi Wang’s laboratory from Ocean University of China (OUC), Dr. Jingjin Xu in Chengtian Zhao’s laboratory from OUC, Dr. Mingjin Xu and Yanan Yin from Qingdao University for providing the experimental materials; Prof. Peter Arvan from University of Michigan Medical School, Dr. Cintia E. Citterio from Chapman University, Dr. Yongxue Li in Zhijun Dong’s laboratory from Yantai Institute of Coastal Zone Research CAS, Prof. Guanglei Liu and Dr. Wenjie Shi from OUC, Ruofei Guo in Hailong Wang’s lab from Shandong University for sharing the experimental methods; Prof. Francesca Coscia from Human Technopole Institute, Prof. Lisui Bao and Runyu Qiao in Shi Wang’s laboratory from OUC for their insight discussion. We thank members of Bo Dong’s laboratory for experimental technical guidance.

This research was funded by the National Key Research and Development Program of China (2022YFC2601302), the Science & Technology Innovation Project of Laoshan Laboratory (No. LSKJ202203204), Shandong Postdoctoral Science Foundation (SDCX-ZG-202400173), and the Natural Science Foundation of Shandong Province (ZR2023MD029), and the Taishan Scholar Program of Shandong Province, China (B.D.).

## Author contributions

Conceptualization: B.D., V.L., J.Z.; Investigation: J.Z., L.K.Y., B.B., Y.M., J.K.W., H.Y.Y.; Analysis and interpretation: J.Z., L.K.Y., B.B.; Funding acquisition: B.D.; Supervision: B.D.; Writing–original draft: J.Z., L.K.Y.; Writing– review and editing: V.L., B.D.

## Additional information

Supplementary materials contain supplementary Figs. 1 to 7, supplementary Tables 1 to 6, and supplementary data 1.

