## Supplementary data for "Identification of the protein precursor for thyroid hormone synthesis in the basal chordate ascidian *Styela clava*"

5. CNRS IRL 2028 "Eco-Evo-Devo of Coral Reef Fish Life Cycle" (EARLY)

6. Institute of Evolution & Marine Biodiversity, Ocean University of China, Qingdao 266003, China.

### These authors contribute equally to this work

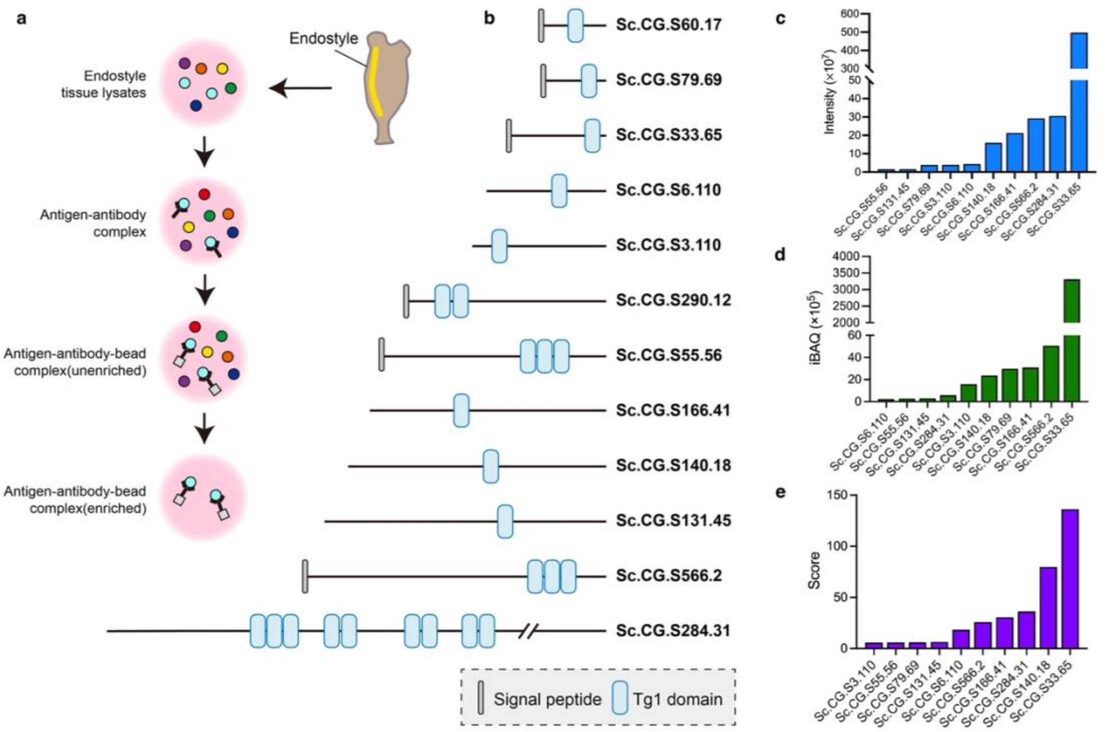


**Supplementary Figure 1. Identification of Thyroglobulin-like (TG-like) proteins in ascidian *Styela clava* through immunoprecipitation (IP). (a)** Workflow of IP experiment for identifying the immunolabeled proteins of antibody of *Bos taurus* TG protein. **(b)** The distributions of signal peptides and Tg1 domains in Tg1-domain-containing proteins in genome of *S. clava*. Signal peptides are indicated in grey, Tg1 domains in blue. **(c-e)** The intensity **(c)**, iBAQ values **(d)** and scores **(e)** of Tg1-domain-containing proteins mapping to the dataset of the immunolabeled proteins of Anti-BtTG.

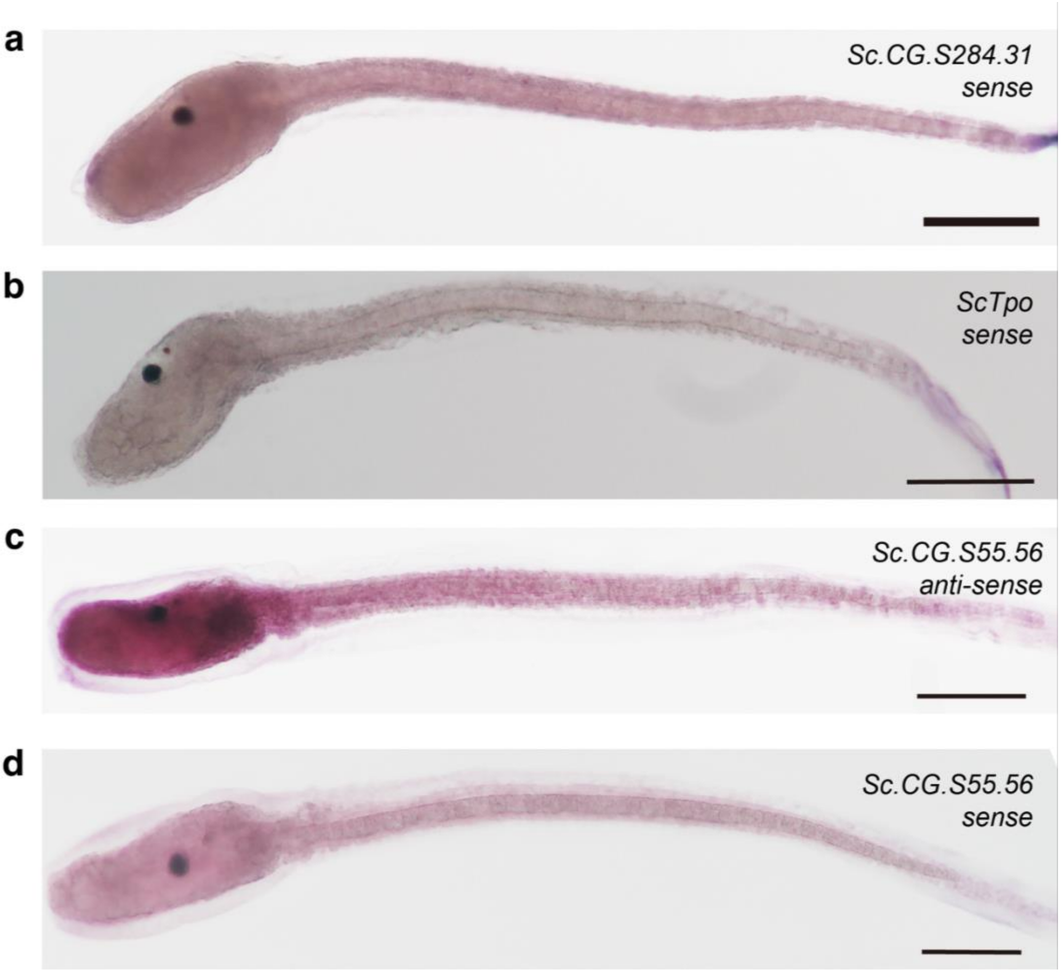


**Supplementary Figure 2. Negative control of whole-mount *in situ* hybridization of *Sc.CG.S284.31* and *ScTpo* genes, and expression patterns of *Sc.CG.S55.56* gene in *S. clava* larvae. (a-d)** *S. clava* larva labeled with sense probes of *Sc.CG.S284.31* gene **(a)** and sense probes of *ScTpo* gene **(b),** anti-sense probes of *Sc.CG.S55.56* gene **(c)** and sense probes of *Sc.CG.S55.56* gene **(d)**. Scale bars = 100 µm.

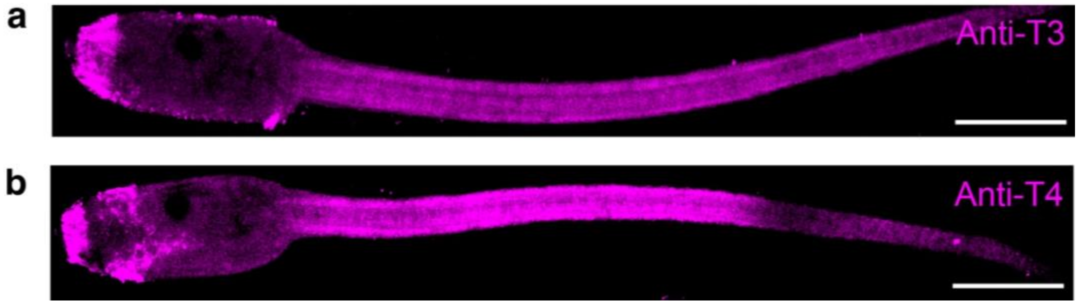


**Supplementary Figure 3. Distributions of THs in *S. clava* larvae. (a-b)** *S. clava* larvae labeled with antibodies of T3 **(a)** and T4 **(b)**. Signals present in magenta. Scale bars = 100 μm.

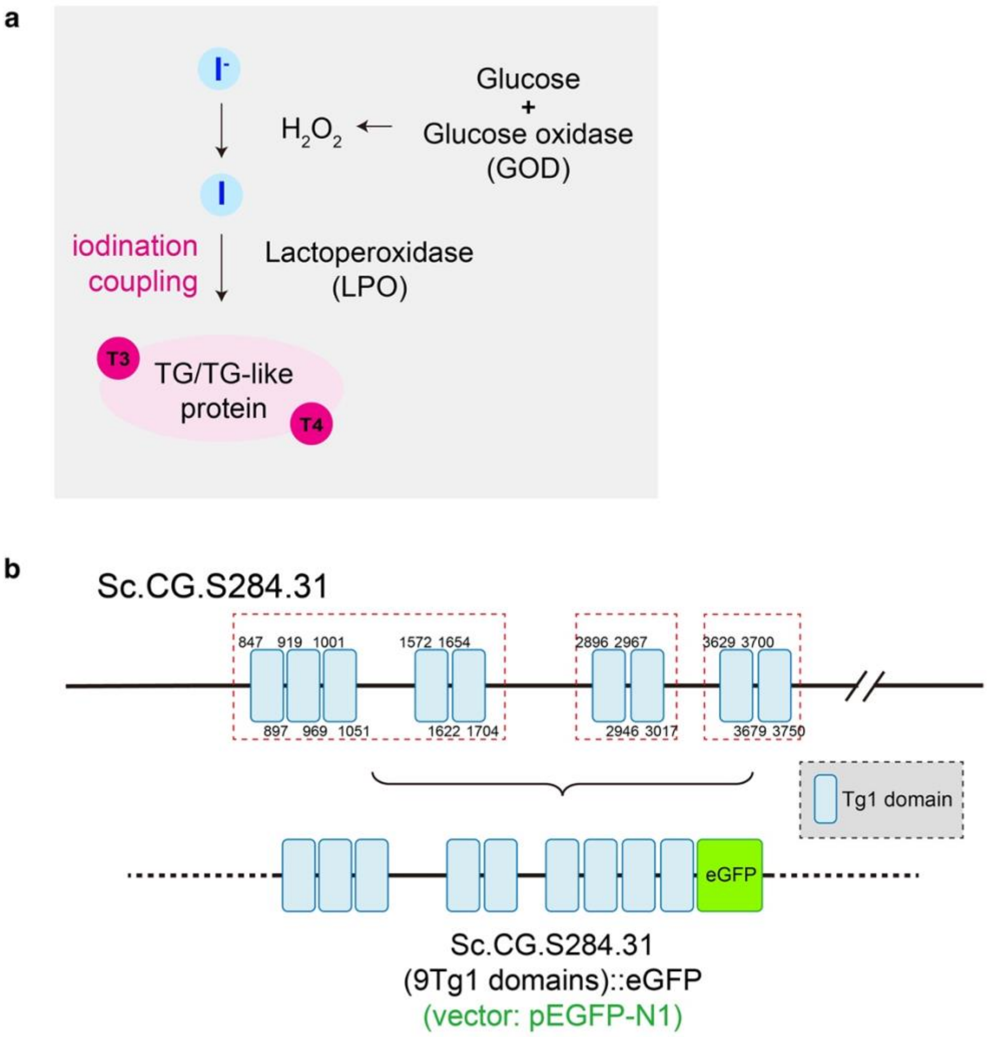


**Supplementary Figure 4. Principle of *in vitro* iodination experiment and schematic diagram of the recombinant Sc.CG.S284.31 protein. (a)** Schematic diagram of *in vitro* iodination experiment. **(b)** Schematic diagram of the recombinant Sc.CG.S284.31 protein. The Tg1 domains are indicated in blue, the eGFP fluorophore in green. The numbers represent the starting (above) and ending (below) positions of the Tg1 domain in the entire amino acid sequence. The amino acid sequences and nucleic acid sequences (including original acid sequences and codon-optimized nucleotide acid sequence) of each Tg1 domains are showed in Supplementary Table 1. The Tg1 domain-enriched fragments of Sc.CG.S284.31 are cloned respectively, and are recombined into the pEGFP-N1 vector (pre-linearization using *NheI* and *KpnI*). The details of plasmid construction are showed in Methods.

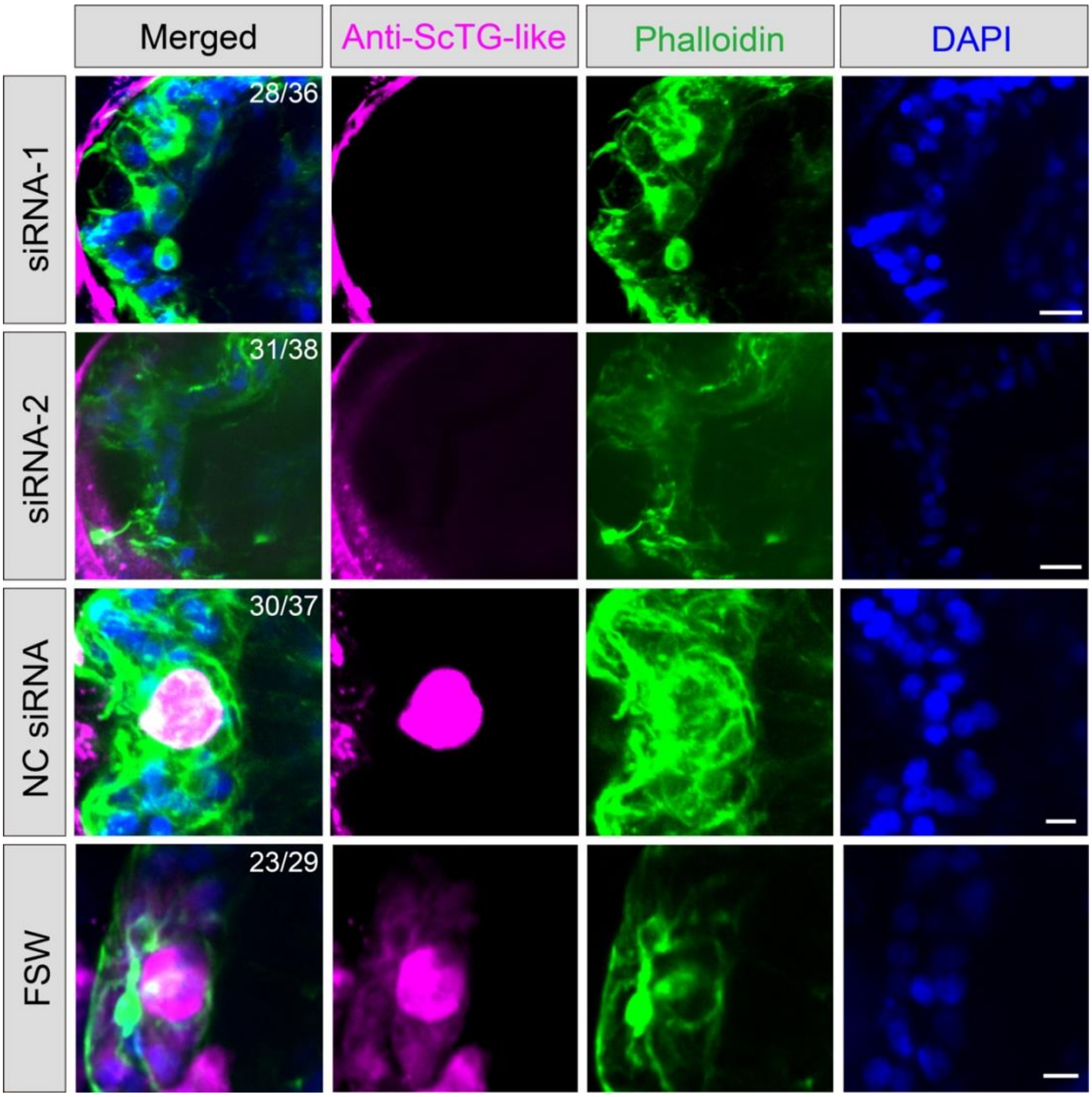


**Supplementary Figure 5. Detection of ScTG-like protein after knocking down the *ScTg-like* gene.** Two specific siRNAs, siRNA-1 and siRNA-2, were used to knock down the expression of *ScTg-like* gene, respectively. Negative control, larvae treated with NC siRNAs. Blank control, larvae treated with FSW. The concentrations of siRNAs are 0.4 µM. The presence of ScTG-like protein in the follicle-like structures was detected using antibody of ScTG-like protein (magenta). The cell boundaries of the follicle-like structures were labeled by phalloidin (green) staining. The nuclei were labeled by DAPI (blue). Scale bars = 10 µm. The fraction showed on each image indicates the number of *S. clava* with the phenotypes showed in the image / total number of *S. clava* in each experimental group.

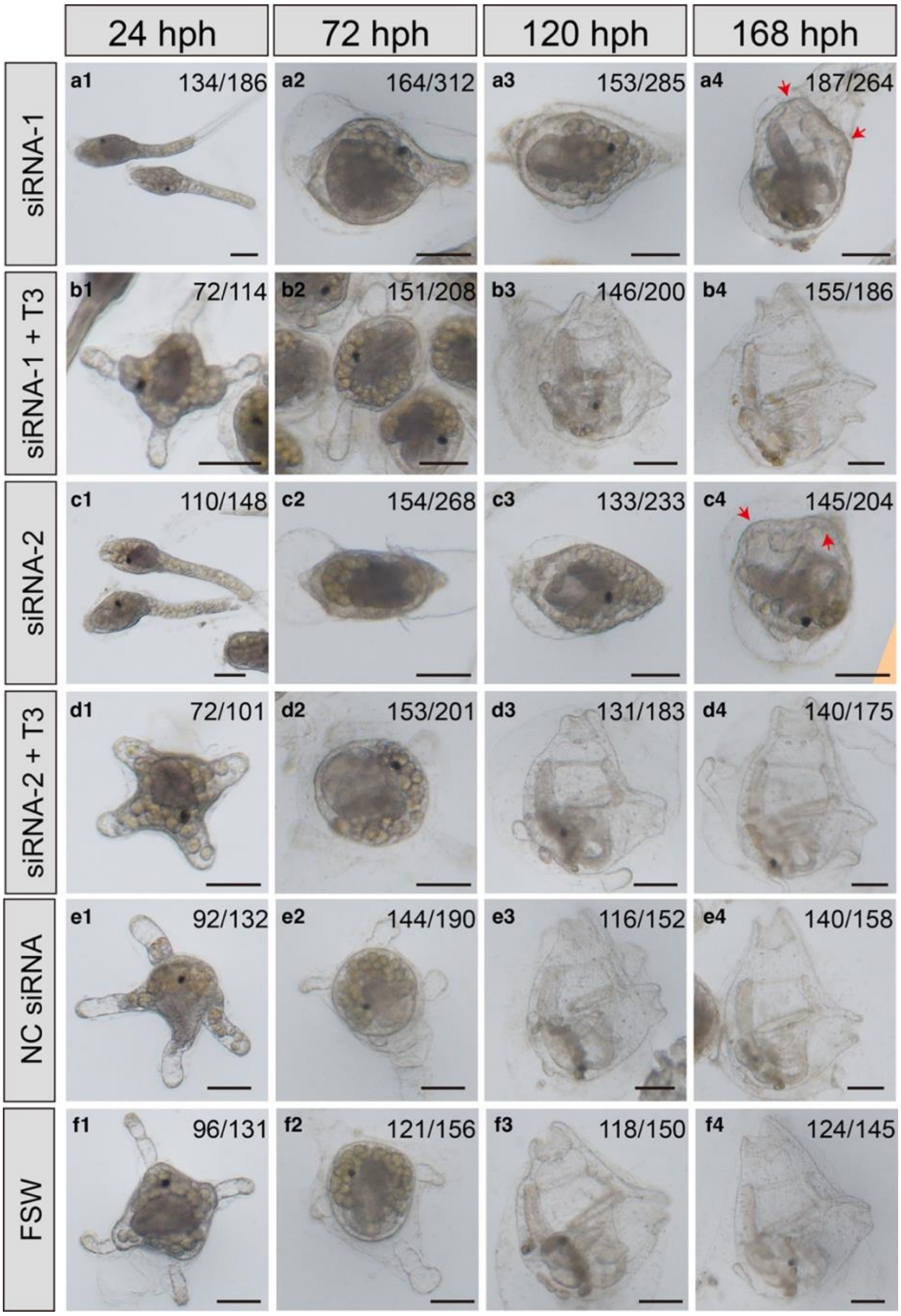


**Supplementary Figure 6. Functional verification of *ScTg-like* in *S. clava* larvae after tail regression.** Two specific siRNAs, siRNA-1 **(a)** and siRNA-2 **(c)**, were used to knock down the expression of *ScTg-like* gene, respectively. Negative control, larvae treated with NC siRNAs **(e)**. Blank control, larvae treated with FSW **(f)**. The concentrations of siRNAs were 0.4 µM. The concentration of MMI was 120 mg/L. T3 was employed for rescue **(b, d)**, with 50 µg/mL. The morphologies of larvae or juveniles were observed at five time points after tail regression, including 24 hours post hatch (hph, **a1-f1**), 72 hph **(a2-f2)**, 120 hph **(a3-f3)** and 168 hph **(a4-f4)**. The red arrows indicate the immature siphons. The fraction showed on each image indicates the number of *S. clava* with the phenotypes showed in the image / total number of *S. clava* in each experimental group. Scale bars = 100 µm.

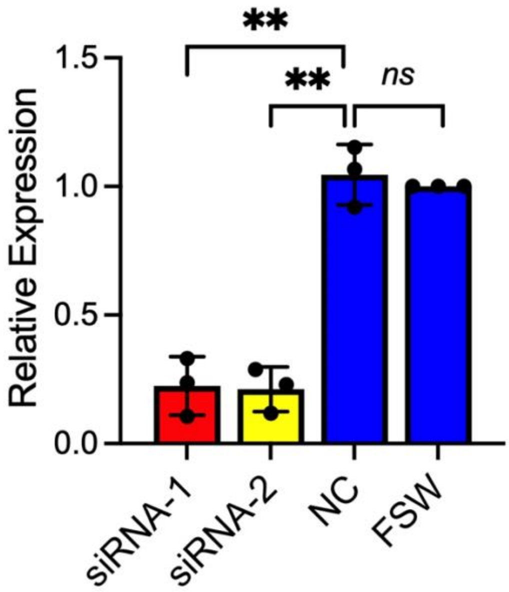


**Supplementary Figure 7. Quantifications of relative expressions of *ScTg-like* gene after treating with siRNAs.** Quantifications of *ScTg-like* mRNA expression in gene knockdown (siRNA-1 and siRNA-2), negative control siRNA treatment (NC), and filtered seawater treatment (FSW) groups. *, $p$-value ≤ 0.05. ns, $p$-value > 0.05, no significant.

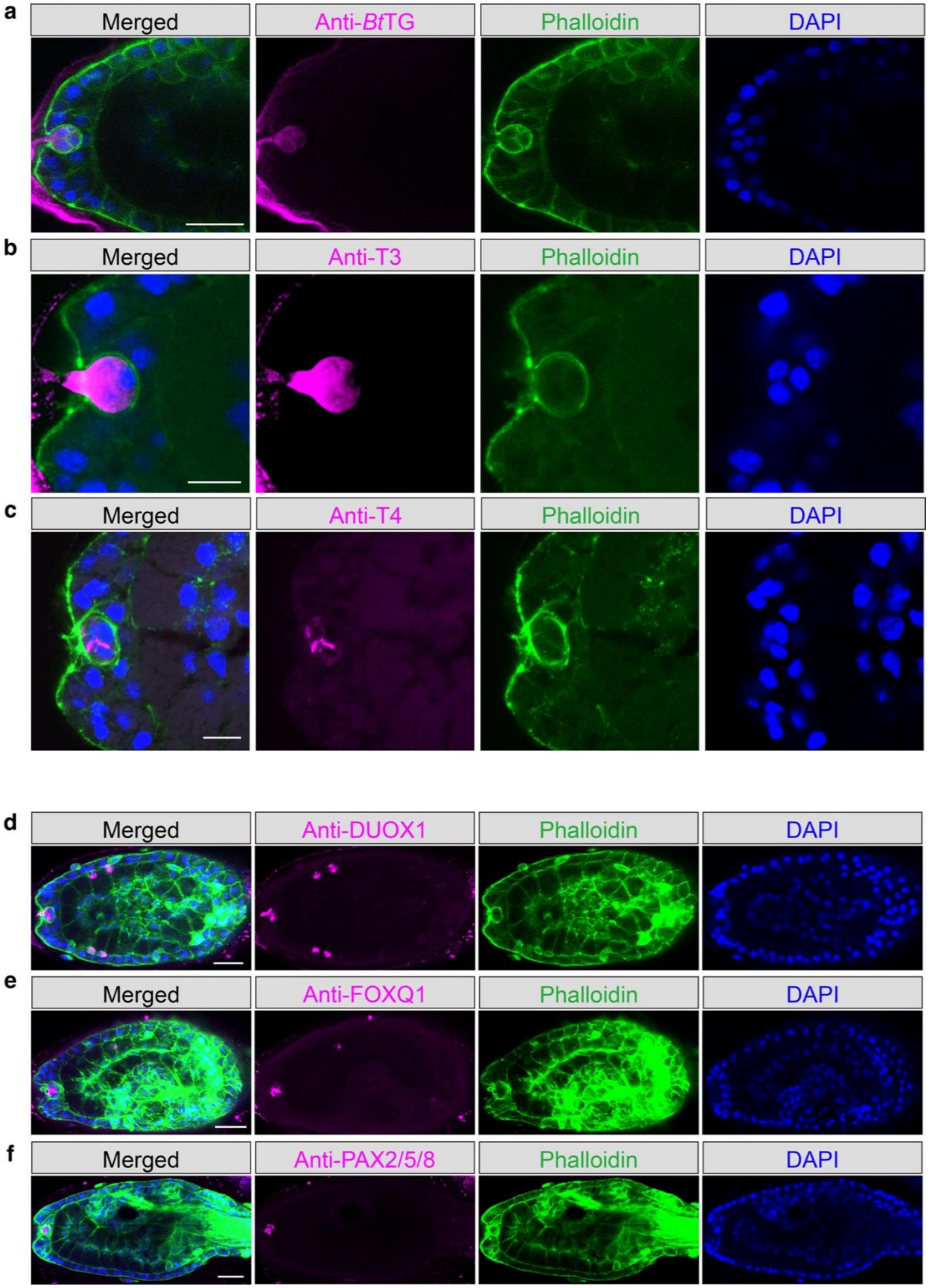


**Supplementary Figure 8. The functions of follicle-like structure are related to TH synthesis and deposition. (a)** The follicle-like structure is labeled by antibody of *Bos taurus* TG protein (magenta), phalloidin (green) and DAPI (blue). Scale bar = 10 µm. **(b-c)** THs are distribution in the follicle-like structures. The follicle-like structures are labeled by antibody of T3 (**b**, magenta) and T4(**c**, magenta), phalloidin (green), and DAPI (blue). Scale bars = 10 µm. **(d-f)** The trunks of *S. clava* larvae are labeled using antibodies of DUOX1 (**d**, magenta), FOXQ1 (**e**, magenta), PAX2/5/8 (**f**, magenta), phalloidin (green), and DAPI (blue). Scale bars = 25 µm.

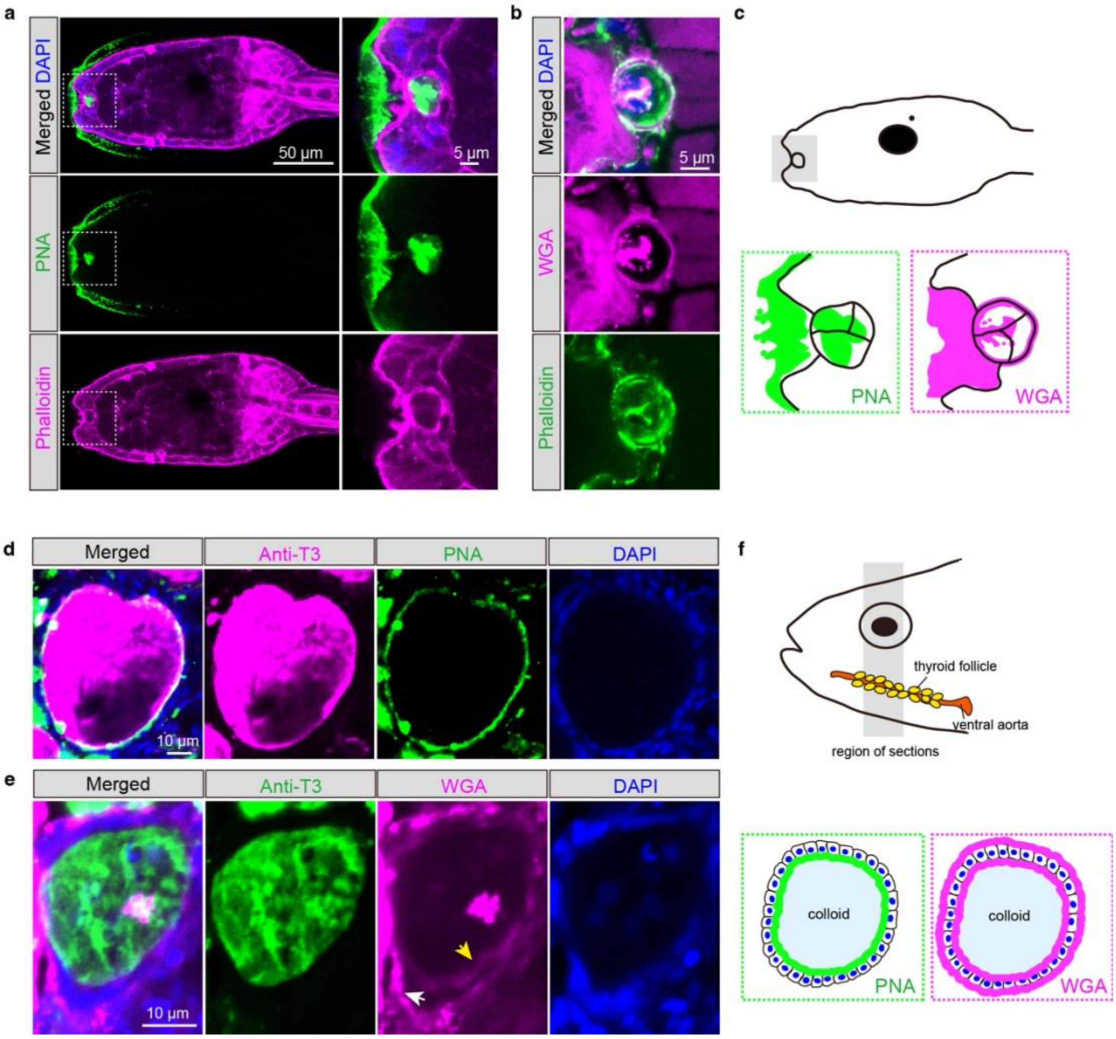


**Supplementary Figure 9. Glycoproteins are enriched in the structures of TH synthesis and storage in both *S. clava* and *Danio rerio*. (a)** The trunks of *S. clava* larvae are labeled using peanut agglutinin (PNA, green), phalloidin (magenta) and DAPI (blue). The whole trunks of *S. clava* larvae are showed on the left in figures, scale bar = 50 µm. The follicle-like structure of each larval is showed on the right, scale bar = 5 µm. **(b)** The follicle-like structure of *S. clava* larvae are labeled using wheat germ agglutinin (WGA, magenta), phalloidin (green) and DAPI (blue). Scale bar = 5 µm. **(c)** Schemes of the results of PNA (green) and WGA (magenta) staining in the follicle-like structure of *S. clava* larvae. **(d)** The thyroid follicle of adult zebrafish *Danio rerio* are labeled using anti-T3 (magenta), PNA (green) and DAPI (blue). Scale bar = 10 µm. **(e)** The thyroid follicle of adult *D. rerio* are labeled using anti-T3 (green), WGA (magenta) and DAPI (blue). The white arrow indicates the basolateral surface of thyroid follicular cells, and the yellow arrow indicates the apical surface. Scale bar = 10 µm. **(f)** Schemes of the results of PNA (green) and WGA (magenta) staining in the thyroid follicle of *D. rerio*.

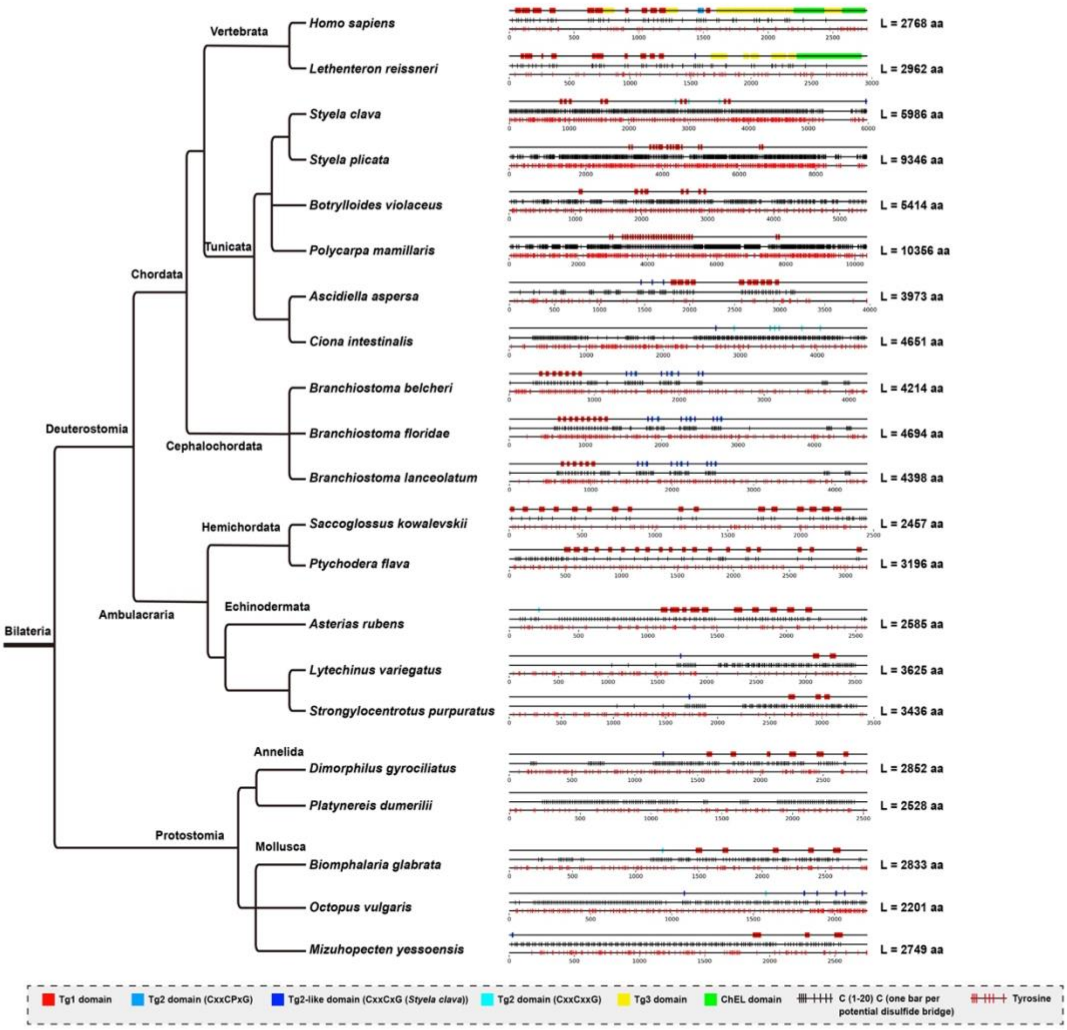


**Supplementary Figure 10. Phylogeny of TG-like candidates in invertebrates and protein architecture of functional domains.** The phylogenetic tree depicts the known relationship between the selected metazoan and illustrates the structural similarity and diversity of TG-like candidates identified by combined Blastp search, specific HMM profiles (Tg1, Tg2, Tg3, ChEL), and conserved regex motifs (disulfide bridges and tyrosine). *Homo sapiens* and *Lethenteron reissneri* represent both extremities of the vertebrate taxon. The *S. clava* sequence is the reference for a functional TG-like protein in invertebrates. The diagrams on the right represent the protein architecture of each candidate, with the position and repetition of Tg1 domains (red), different types of Tg2 domains (including Tg2 domain (C××CP×G) in blue, Tg2-like domain (C××C×G) in dark blue, Tg2-like domain (C××C××G) in light blue), Tg3 (yellow) and cholinesterase-like (ChEL) domain (green). The horizontal black bars below the domains indicate potential disulfide bonds (one per pair of cysteines spaced 1 to 20 amino acids apart), essential for structural stability. The red lines at the bottom correspond to tyrosine residues. This combined representation allows the identification of conserved protein architectures and potential functional similarities. The length of TG-like candidate is labelled on the right. The information of each protein is listed in Supplementary Table 4.

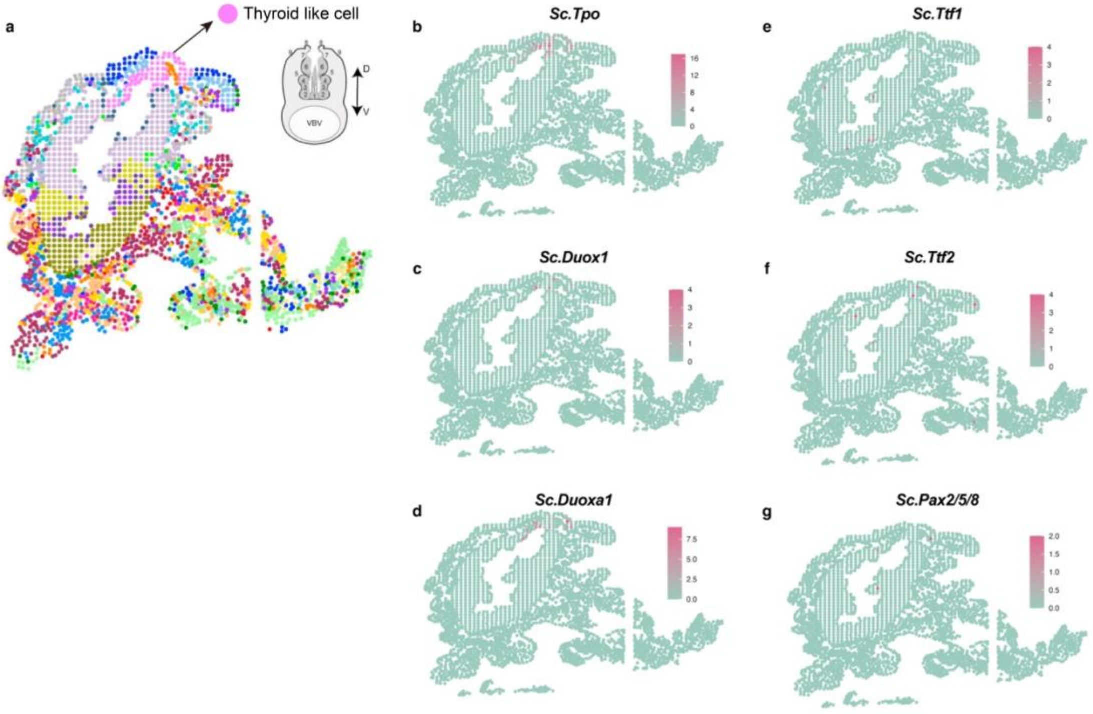


**Supplementary Figure 11. Distributions of crucial TH synthesis-related genes in adult endostyle. (a)** Spatial atlas shows the distribution of cell units of *S. clava* endostyle on the transverse section colored by cell annotation. The thyroid-like cells in *S. clava* endostyle are labelled by pink dots. The Distribution of (b) *Sc.Tpo*, (c) *Sc.Duox1*, (d) *Sc.Duoxa1*, (e) *Sc.Ttf1*, (f) *Sc.Ttf2* and (g) *Sc.Pax2/5/8* genes in *S. clava* endostyle based on the spatial single cell transcriptomic data. Higer expression level is showed in pink, and the lower expression level is showed in green.
